# Contrasting patterns of spatial genetic structure in endangered southern damselfly (*Coenagrion mercuriale*) populations facing habitat fragmentation and urbanisation

**DOI:** 10.1101/2024.05.02.592171

**Authors:** Agathe Lévêque, Anne Duputié, Vincent Vignon, Fabien Duez, Cécile Godé, Cédric Vanappelghem, Jean-François Arnaud

**Affiliations:** Univ. Lille, CNRS, UMR 8198 - Evo-Eco-Paleo, F-59000 Lille, France; Office de Génie Ecologique (O.G.E.), F-67200 Strasbourg, France; Office de Génie Ecologique (O.G.E.), F-94100 Saint-Maur-des-Fossés, France; Conservatoire d’espaces naturels des Hauts-de-France, F-80480 Dury, France

**Keywords:** central-marginal hypothesis, conservation genetics, central and peripheral populations, genetic diversity, isolation-by-distance, population genetic structure, urbanisation effects

## Abstract

**Aim:** Human-induced environmental changes result in habitat loss and fragmentation, impacting wildlife population genetic structure and evolution. Urbanised and geographically peripheral areas often represent unfavourable environments, reducing connectivity among populations and causing higher population genetic differentiation and lower intra-population genetic diversity. We examined how geographic peripherality and anthropogenic pressures affect genetic diversity and genetic differentiation in the protected southern damselfly (*Coenagrion mercuriale*, Odonata), which has low dispersal capabilities and specific habitat requirements and whose populations are declining.

**Location:** We studied two areas: one in semi-natural habitats at the periphery of the species geographic range (northern France) and the other more central to the species’ range, in an urbanised area surrounding the city of Strasbourg (Alsace, eastern France).

**Methods:** We genotyped 2743 individuals from 128 populations using eleven microsatellite loci. We analysed the spatial distribution of neutral genetic diversity (allelic richness, heterozygosity, levels of inbreeding, genetic relatedness), the extent of genetic differentiation, and population affiliations (sPCA analyses) within the two areas. We also examined fine-scale patterns of gene flow in the urbanised area of Alsace by investigating patterns of isolation by distance and estimating effective migration surfaces (EEMS method).

**Results:** Northern peripheral populations showed lower levels of genetic diversity and higher levels of genetic differentiation than central Alsacian populations. Although located in anthropised habitats, geographically central Alsacian populations showed high levels of gene flow, with dispersal events mainly occurring overland and not restricted to watercourses. However, the highly urbanised city of Strasbourg negatively impacted nearby populations by reducing levels of genetic diversity and increasing population genetic differentiation.

**Main conclusions:** These results showed the need for management action by restoring breeding sites and creating migratory corridors for peripheral southern damselfly populations. However, our results also highlighted the resilience of southern damselfly in central range populations facing strong urbanisation pressures.

## Introduction

In the context of major environmental changes caused by human activities, such as urbanisation, intensive agriculture or land-use changes, habitat loss and fragmentation are major threats to biodiversity (Fahrig, 2003; Wilson *et al*., 2016). Fragmented populations face a reduction both in the amount of favourable habitats and in gene flow among populations inhabiting the remaining habitat patches (Fahrig, 2003). This may generate both demographic and genetic effects on populations, with an increased risk of inbreeding, a loss of genetic diversity, and an increase in genetic differentiation among populations, which can ultimately lead to a vortex of extinction in the long term (Frankham *et al*., 2013; Ellegren and Galtier, 2016).

Urbanisation also impacts the fate of wildlife populations, as described by the “urban fragmentation” model in which habitat fragmentation and degradation magnify the extent of genetic drift and decrease gene flow events among populations (Johnson and Munshi-South, 2017; Miles *et al*., 2019). This process was indeed documented in numerous species of different taxa (Desender *et al*., 2005; Vandergast *et al*., 2006; Munshi-South *et al*., 2016; Miles *et al*., 2019; Trumbo *et al*., 2019; Fusco *et al*., 2021). Nonetheless, the impact of urbanisation varies greatly, depending on the species’ life-history traits but also on the landscape matrix (Miles *et al*., 2019; Kimmig *et al*., 2020; Richardson *et al*., 2021). Urbanisation can not only reduce connectivity between populations, but also create barriers or corridors that influence species’ dispersal behaviours and pathways (Kimmig *et al*., 2020). While highly mobile species are more likely to maintain some connectivity with exurban populations, avoiding inbreeding depression and preserving high levels of genetic diversity, this might not be the case for low vagility species depending on specific habitats and facing dispersal barriers (Fusco *et al*., 2021; Johnson & Munshi-South, 2017; Richardson *et al*., 2021).

Yet, to obtain a global vision of the conservation management of a species, it is important to understand patterns of spatial genetic structure both at local scales and at the larger scale of the species’ geographical range (Guo, 2012; DeWoody *et al*., 2021; Willi *et al*., 2022). The “abundant centre” model posits that populations near the centre of a species’ geographic distribution are the most abundant, and become smaller and scarcer toward the periphery of the range (Brown, 1984). Two main predictions arise from this model: (i) peripheral populations should exhibit lower levels of genetic diversity compared to central ones, as a result of genetic drift associated with small population size; (ii) peripheral populations are likely to exhibit higher levels of genetic differentiation than central populations, because of genetic drift and of restricted gene flow due to geographic isolation (Vucetich and Waite, 2003; Eckert *et al*., 2008). Nonetheless, this pattern is not systematically observed, as the geometry of the species’ range, historical and current ecological factors can lead to deviations from this model (Eckert *et al*., 2008; Guo, 2012; Johansson *et al*., 2013; Pironon *et al*., 2017).

All these evolutionary considerations are particularly relevant to the Odonata taxa, since they are bioindicator species for freshwater ecosystems and have a biphasic life cycle, with aquatic larval and terrestrial adult stages, making them dependent on both environments for their development. While Odonata species are considered to be efficient fliers, potentially able to disperse over large spatial distances, some Zygoptera species, such as the southern damselfly (*Coenagrion mercuriale,* Charpentier, 1840), have low dispersal capability. Mark-recapture studies have documented rare long-distance movements of southern damselflies over more than one kilometre (Purse *et al*., 2003; Rouquette and Thompson, 2007; Watts *et al*., 2004c). The species is geographically restricted to western Europe, with northern range limits in the UK, France, and Belgium, eastern limit in Germany and southern limit in northern Africa (Grand, 1996; see Figure S1). Southern damselflies are under high conservation priority because the species has become almost extinct in seven European countries on the northern and eastern boundaries of its distribution, where many populations are threatened by the deterioration and loss of its habitat associated with changes in agricultural practices (Grand, 1996). The southern damselfly is found in lotic habitats, requiring slow-flowing small streams or ditches, open habitats with little shade, and the presence of helophytes (Rouquette and Thompson, 2005; Purse and Thompson, 2009).

In this study, using a set of microsatellite loci, we aimed to examine the spatial patterns of genetic diversity and genetic differentiation within and among southern damselfly populations in two contrasting areas: (i) an urbanised area located around the city of Strasbourg in Alsace (eastern France), towards the centre of the species’ geographic distribution and characterised by the occurrence of numerous populations; (ii) a second area located in northern France and Belgium, towards the northern limit of the species’ distribution, where populations are scarcer and only occur in semi-natural habitats. We asked the following questions:

1. Do central urbanised and peripheral semi-natural populations of southern damselfly exhibit the same levels of genetic diversity and genetic differentiation? We expected populations in northern France and Belgium to show higher levels of genetic differentiation and lower levels of genetic diversity than populations in Alsace, due to their geographical isolation close to the limit of the species’ distribution. However, these differences could be mitigated by a negative effect of urbanisation on Alsacian populations, even though they are more central to the species’ range.
2. When focussing on the Alsace region, where populations are densely distributed, do migration pathways occur overland or through the hydrographic network? We compared the patterns of isolation-by-distance (IBD) using ecological distances measured along watercourses to those found using simple straight-line Euclidean geographic distances. We only conducted this analysis in Alsace, because of a larger occurrence of the species and, consequently, because of a larger number of sampled populations.
3. Does the urban pressure of the city of Strasbourg impact population genetic features of neighbouring southern damselfly populations? The main expectation was that the city represents a substantial barrier to the dispersal of southern damselflies. This could lead to a decrease in levels of intra-population genetic diversity for populations close to the city, according to the urban fragmentation model (Miles *et al*., 2019).

## Methods

### Study sites and sample collection

We sampled two areas: (i) Northern France and Southern Belgium (Figures 1A-C); (ii) the Alsace region, located in eastern France (Figures 1B-C).

**Figure 1:**
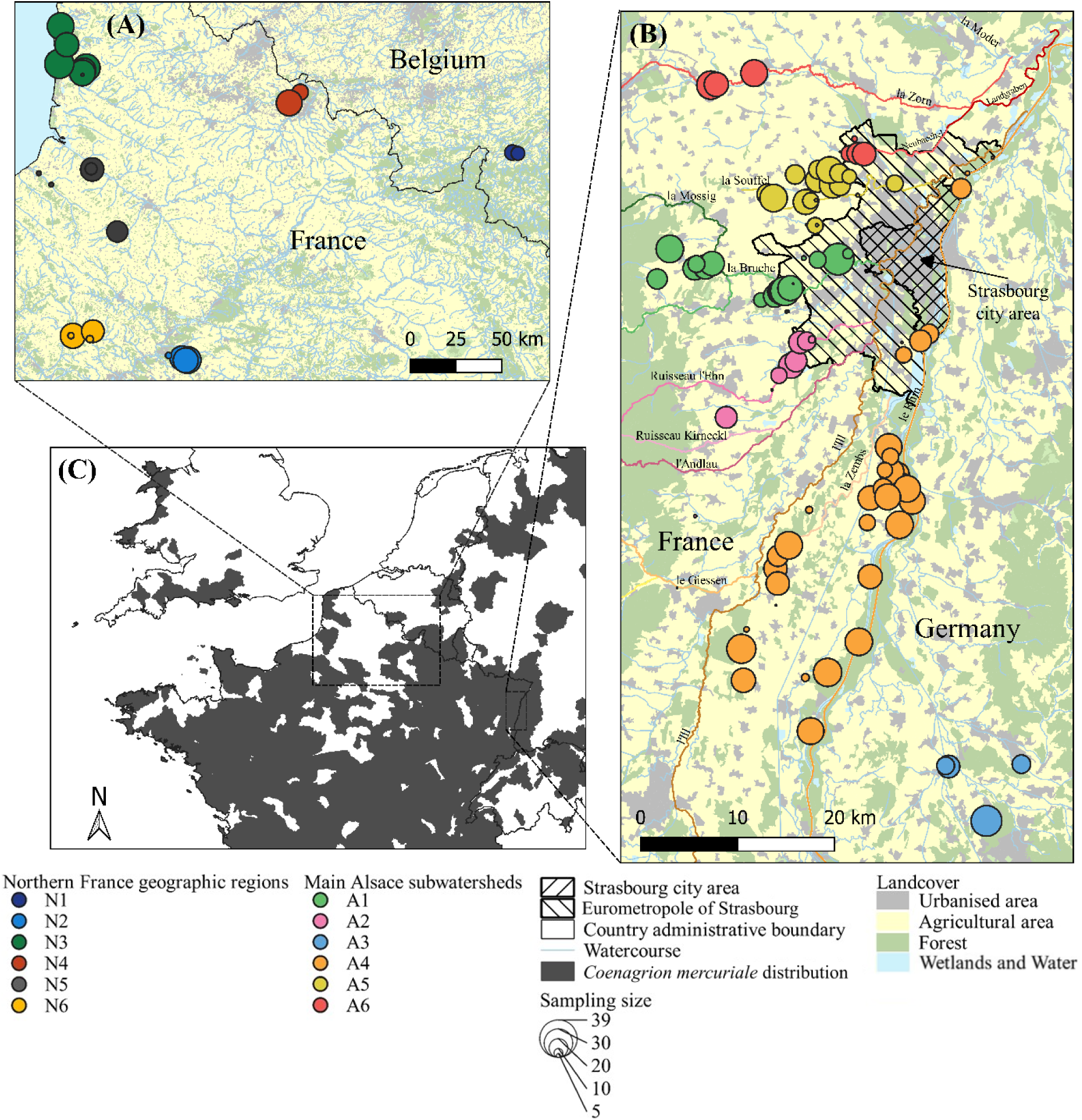
Location of 128 populations of southern damselfly (*Coenagrion mercuriale*) within two study areas: Northern France **(A)**, and Alsace **(B)**. Each circle represents one population, with circle size being proportional to the number of individuals sampled. Strasbourg city and the eurometropole of Strasbourg are represented by the striped areas. Around Strasbourg, main watercourses are represented and coloured depending on the hydrographic network they contribute to. Circle colours correspond to the main subwatershed or geographical region each population belongs to. Land cover was simplified from Corine Land Cover Edition 2018. **(C)** The grey zones represent the current geographic distribution of *Coenagrion mercuriale* (data obtained from IUCN SSC Odonata Specialist Group 2019. The IUCN Red List of Threatened Species. Version 2022-2).

In Northern France and southern Belgium, southern damselfly populations are scarce (Figure S1). We thus considered this region to be at the northern limit of the species’ geographic distribution. This region is very urbanised with intensive agriculture, and is characterised by a low occurrence of natural areas. In this region, southern damselflies are only found in semi-natural sites (Fierimonte and Vanappelchem, 2021). The “Conservatoire des Espaces Naturels Hauts de France” (CEN) is in charge of managing biodiversity in more than 540 semi-natural sites in northern France, and surveys additional sites when required. Because southern damselfly is an endangered species in this region, its occurrence is well recorded by biodiversity managers (see Fig. S2A). Over this wide region (∼60 000 km²), we exhaustively sampled all 24 semi-natural areas where southern damselfly was documented by the CEN (Fierimonte and Vanappelchem, 2021). These sites are located in six subregions labelled N1 to N6 (Figure 1A; N1: Belgium; N2: Oise; N3: Opal coast; N4: Scarpe; N5: Somme; N6: Vexin).

In Alsace region (and in neighbouring areas), southern damselfly populations are more densely distributed (Figures S1, S2B). We thus considered the Alsace region to be more central to the species’ geographic distribution. In this region, southern damselflies can occur in natural or in more anthropised environments. We thus surveyed an area encompassing the city of Strasbourg. This city spans 78 km² and counts about 290 000 inhabitants, with a human population density of more than 3700 people/km². Strasbourg city is located within the urban area named “Eurométropole de Strasbourg” with about 500 000 inhabitants for 330 km². A few kilometres west are the Vosges mountains, where southern damselflies do not occur. In this region, the landscape is characterised by intensive and less intensive agriculture (mostly cereals and grapevine on the slopes). For this region of about 4 000 km², we accessed to historical records of southern damselfly occurrence (http://association.imago.free.fr/; https://www.gbif.org/fr/species/1422012). We therefore surveyed all known sites where this species was reported, but we also investigated additional sites that appeared to present favourable conditions for southern damselflies, by selecting them on Google Earth, along small streams outside forest areas. In total, we surveyed 206 sites in Alsace (see Fig. S2B), and were able to find southern damselfly populations in 104 of them. Populations were then spatially grouped into six main subwatersheds named A1 to A6 (Figure 1B; A1: Bruche/Mossig; A2: Ehn/Andlau; A3: Dreisan/Elz; A4: Ill/Rhine; A5: Souffel; A6: Zorn/Landgraben).

We collected adult individuals of southern damselfly (*Coenagrion mercuriale*) using an insect net in spring (May to July) from 2017 to 2022 for Northern France and from 2021 to 2022 for the Alsace sampling area. Given the limited dispersal capability of the southern damselfly (Watts *et al*., 2004c; 2006), we designed sampling sites following 200 metres sections located along rivers and streams, separated by at least 500 metres to avoid repeated sampling of the same population (Figure S2B).

We sampled a total of 2743 adult specimens (545 in Northern France, 2198 in Alsace; 1-39 per population; mean 21.4 ± 11.3; Figure 1 and Table S1), with a sex ratio strongly biased towards males (86.73%) due to their higher-flying activities and conspicuousness. Table S1 lists the geographic coordinates and sampling sizes of each population. We collected two kinds of samples: whole individual body, or only the right middle leg of each individual, the latter method being non-lethal and does not impact the damselfly’s survival (Fincke and Hadrys, 2001). All samples were stored in 100% ethanol until DNA extraction.

### Genotyping

For whole individuals, we removed abdomens from the rest of the body to avoid sampling intestinal microbiota, crushed the samples using five MN Beads Type D (Macherey-Nagel). We crushed leg samples with only three beads. We extracted total genomic DNA from each whole individual sample using NucleoMag® Tissue Kit (Macherey-Nagel) according to the manufacturer’s recommendations. For leg samples, we purified DNA with 12µL NucleoMag B-beads and 12µL of pure water, and eluted in 50 µL.

We genotyped all samples using eleven unlinked nuclear microsatellite loci named LIST002, LIST023, LIST034, LIST035, LIST037, LIST042, LIST062, LIST024, LIST060, LIST063, and LIST066, isolated and described in Watts *et al* (2004a,b). Locus LIST060 was excluded from all subsequent analyses because it showed evidences for null alleles occurrence (*F*IS = 0.377, *P* < 10^-3^; Table S2). PCR reactions were performed in two multiplexes described in Table S2. PCR reactions were conducted in a volume of 10 µL, using 3 µL of DNA (0.5-5 ng/mL), 5 µL Multiplex PCR Master Mix (QIAGEN) and a primer mix (each primer at a final concentration of 0.2 µM). PCR amplifications were as follows: (i) 15 min at 95°C, (ii) 30 cycles for whole individuals, and 32 cycles for legs of 30 s denaturation at 94°C, 90 s annealing 55°C, and 60 s elongation at 72°C, (iii) 30 min at 60°C. 1.5 µL of PCR products (1/10 diluted for whole individual samples) were pooled with 0.25 μL of RadiantDy^TM^ 632 500 MOB size standard (Eurogentec, Seraing, Belgium) and 9.75 μL of formamide (Applied Biosystems, Foster City, CA), electrophoresed and sized with an ABI PRISM 3130XL sequencer (Applied Biosystems) and the software GeneMapper 5.0 software (Applied Biosystems). Amplification failure was scarce (0% for Alsace and 0.14% for Northern France).

### Genetic diversity

Standard population genetic statistics were calculated using FSTAT 2.9.4 (Goudet, 2003) and R (version 4.2.1; R Core Team, 2022) libraries ‘adegenet’ 2.1.9 (Jombart, 2008), ‘hierfstat’ 0.5-11 (Goudet, 2005), ‘pegas’ 1.1 (Paradis, 2010), and ‘demerelate’ 0.9 (Kraemer and Gerlach, 2017). All statistics were weighted by population sizes (Weir and Cockerham, 1984; Nei, 1987), allowing for non-biased comparisons across populations and across regions. Moreover, to ensure statistically robust results, we estimated these standard statistics of genetic diversity for only populations with a minimal sampling size of at least eight individuals, allowing to compare relevant genetic diversity estimates between representative samples of populations (El Mousadik and Petit, 1996; Hedrick, 2011; Frankham *et al*., 2013). These basic statistics included the total number of alleles sampled (*A*T), the allelic richness (*A*r), the observed (*H*o) and the expected (*H*e) heterozygosity, and the mean individual kinship coefficient within populations (*F*ij; Loiselle *et al*., 1995). Statistical differences between Alsace and Northern France in terms of mean levels of genetic diversity and of mean levels of intra-population kinship were tested by permutation tests using the R library ‘perm’ 1.0-0.4 (Fay and Shaw, 2010).

To investigate population genetic structure over the whole dataset and among the two study areas and the different subwatersheds and subregions, we computed estimates of *F*-statistics following Weir and Cockerham (1984) and tested their statistical significance using permutation tests (10000 permutations) implemented in FSTAT (Goudet, 2003). As for the calculation of basic genetic diversity estimates, *F*-statistics estimates were only used for populations with at least eight individuals.

To study the potential effect of Strasbourg city on southern damselfly populations, we considered a specific radius of 20 km around the city centre and we performed a linear regression between the distance of each population from the centre of Strasbourg city and i) the genetic diversity estimates (*A*r, *H*e), ii) the mean individual kinship coefficient within populations (*F*ij) and iii) the mean pairwise levels of *F*ST calculated according to Weir and Cockerham (1984) between each focal population and all other populations within this radius. Because other ecological factors, such as land-use, forest occurrence or the presence of the Vosges mountains are expected to shape the genetic structure at larger scales of investigation, we restricted this analysis to a 20 km radius around the centre of Strasbourg to avoid confounding effects (Figure S3).

### Delimitation of population boundaries and genetic discontinuities

Genetic discontinuities and grouping of genetically related populations were assessed using spatial Principal Component Analysis (sPCA). This spatially explicit multivariate method, implemented in the ‘adegenet’ R library (Jombart, 2008), reveals spatial genetic patterns and detects cryptic geographical variation in allele frequencies allowing to depict population boundaries, without making assumptions with regard to linkage disequilibrium or departures from Hardy-Weinberg equilibrium (Jombart *et al*., 2008). We used the Delaunay triangulation graph to create the spatial network underlying the sPCA. To draw a comprehensive synthetic representation, we represented population coordinates along the first three principal components on the Red, Green, and Blue colour channels as in Menozzi *et al* (1978).

### Analysis of spatial genetic structure

We did not investigate the patterns of isolation-by-distance (IBD) in northern France, both because of the smaller number of sampled populations in this area and because of the difference in geographic scale of sampling. Indeed, because southern damselflies have limited dispersal capabilities, it is not expected to find a migration/drift equilibrium among distant and isolated subregions. Therefore, we studied the fine-scaled spatial genetic structure by only focusing on the whole Alsace region and in the three major subwatersheds, where substantial numbers of populations and individuals ensure biologically relevant results. To search for an IBD pattern expected under migration/drift equilibrium, we considered two different estimations of geographical distances: (i) log-transformed Euclidean distances (dEucli); (ii) log-transformed shortest path distances along watercourses (dStream) because southern damselflies only reproduce along watercourses. We computed Euclidean geographical distances using the R function *dist*, and geographical distances along waterways using the R package ‘riverdist’ 0.15.5 (Tyers, 2022). To calculate the latter distance, we used the BD TOPAGE® 2019 linear hydrology shapefile. We then assessed the spatial genetic structure using two complementary approaches.

First, to test for IBD among populations, we regressed pairwise *F*ST estimates against geographical distances and tested the significance of the Mantel statistic *rz* (Smouse *et al*., 1986) using the *mantel.rtest* function of the R package ‘ade4’ with 10000 permutations (Dray and Dufour, 2007).

Secondly, to get insight into the spatial genetic structure over the Alsace area, we analysed the relationship between pairwise individual kinship coefficients *F*ij (Loiselle *et al*., 1995) and geographical distance using SPAGEDI 1.5 (Hardy and Vekemans, 2002). To allow comparable and relevant statistical results in terms of mean genetic relatedness among individuals, nine classes of increasing geographical distances were defined, which included nearly identical number (mean of 268278 per distance class) of pairwise individual comparisons. In each distance class, we assessed 95% upper and lower confidence intervals of *F*ij using 10000 permutations of individual locations. Finally, to compare the strength of spatial genetic structure between the different geographical distances and subwatersheds in Alsace, we used the *Sp* statistic (Vekemans and Hardy, 2004).

### Estimated Effective Migration Surface (EEMS)

Finally, to identify geographical regions where migration rate was higher or lower than expected under an isolation-by-distance null model, we used the Estimated Effective Migration Surfaces method (EEMS; Petkova *et al*., 2016) in the whole Alsace region. We ran the model for a range of deme values (400, 1000, 1500, and 2000) and parameters were optimised until the proposals were accepted about ∼20-30% of the time, as recommended by Petkova *et al*. (2016). Convergence of MCMC chains, checked using the ‘reemsplots2’ R library (Petkova, 2022), was reached with the MCMC parameters by default (numMCMCIter = 2000000, numBurnIter = 1000000, numThinIter = 9999). This analysis was again restricted to the Alsace region, where populations were more densely distributed.

## Results

### Genetic diversity

Over all populations, single-locus *F*IS values did not significantly differ from zero, except for locus LIST037 (*F*IS = 0.035, *P* < 0.05; Table S2). Single-locus *F*ST all significantly differed from zero, with a mean multilocus estimate of 0.063 (*P <* 0.05; Table S2). The number of alleles per locus ranged from two to 24, the mean *H*e was 0.568, and the mean *H*o was 0.507 (Table S2). Within populations, we detected no significant departures from Hardy-Weinberg equilibrium (Table S1).

In Alsace, levels of genetic diversity were high, with a mean allelic richness of 3.19 ± 0.13, a mean *H*e of 0.529 ± 0.021 and a mean *H*o of 0.528 ± 0.03 (Table S1, Figure 2A, Figure S4). The geographical patterns of these estimates of genetic diversity are displayed in Figure S5. The mean intra-population kinship *F*ij was 0.022 (Table S1, Figure 2B). Five populations showed higher mean levels of genetic relatedness between individuals than other Alsacian population (populations Gox1, S17, S21, Rlei45, Rsc1; Table S1, locations visible on Figure S3). Gene diversity (*H*e) and allelic richness (*A*r) increased with increasing distance of populations from the centre of Strasbourg within a radius of 20 km around the city (Figure 3A, Figure S6A). On the contrary, mean pairwise *F*ST levels and mean intra-population kinship *F*ij decreased with increasing distance of populations from the centre of Strasbourg (Figures 3B, S6B). This significant relationship observed between genetic diversity or genetic differentiation and the geographical distance from the city of Strasbourg disappear beyond this scale of 20 km, which suggested other ecological factors than strict urbanisation level impacting population genetic features at larger scale of observation (data not shown).

**Figure 2:**
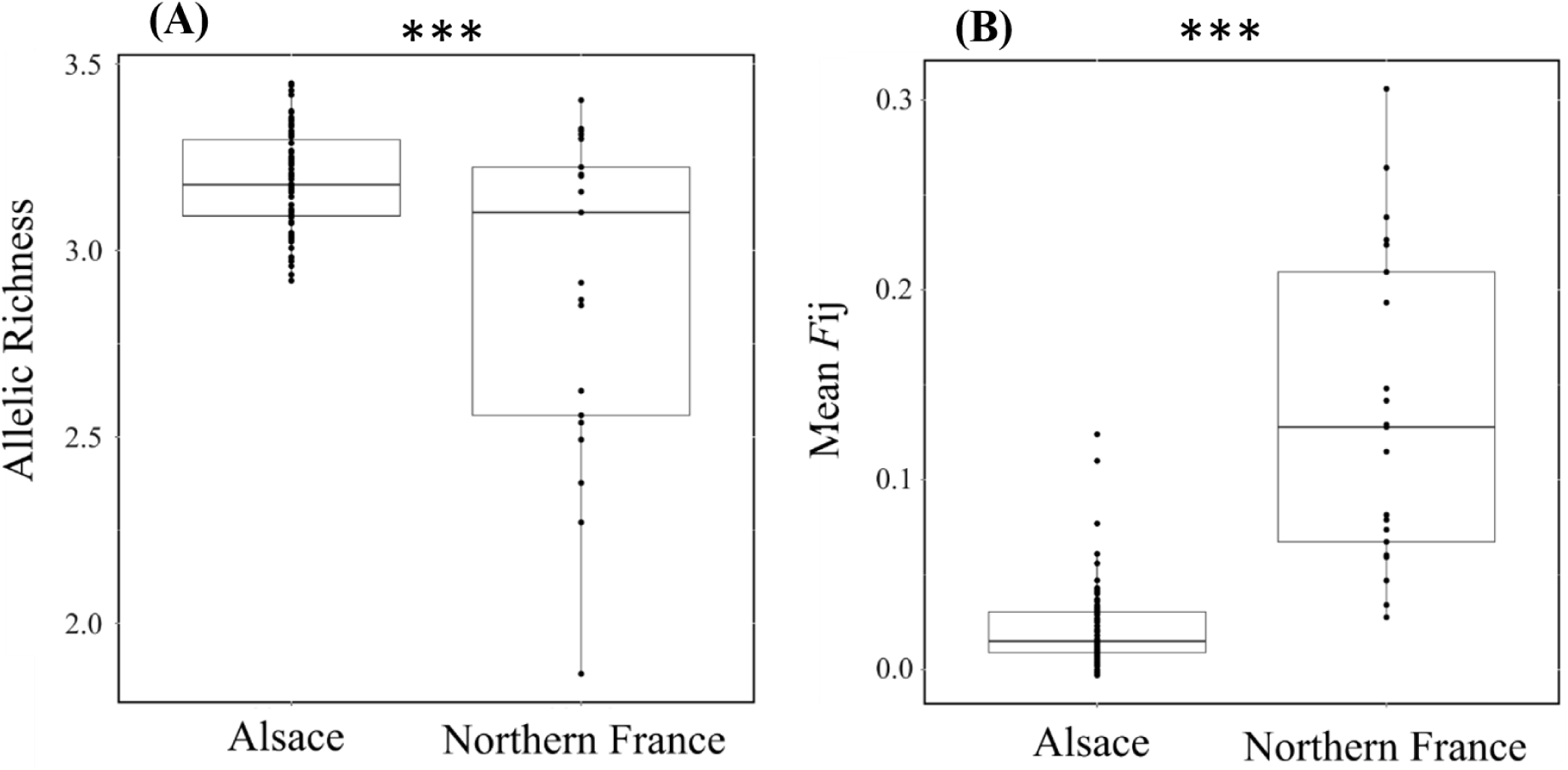
Distribution of allelic richness *A*r **(A)** and mean kinship coefficient *F*ij **(B)** in the southern damselfly within the Alsace and Northern France areas. ***: *P* < 0.001.

**Figure 3:**
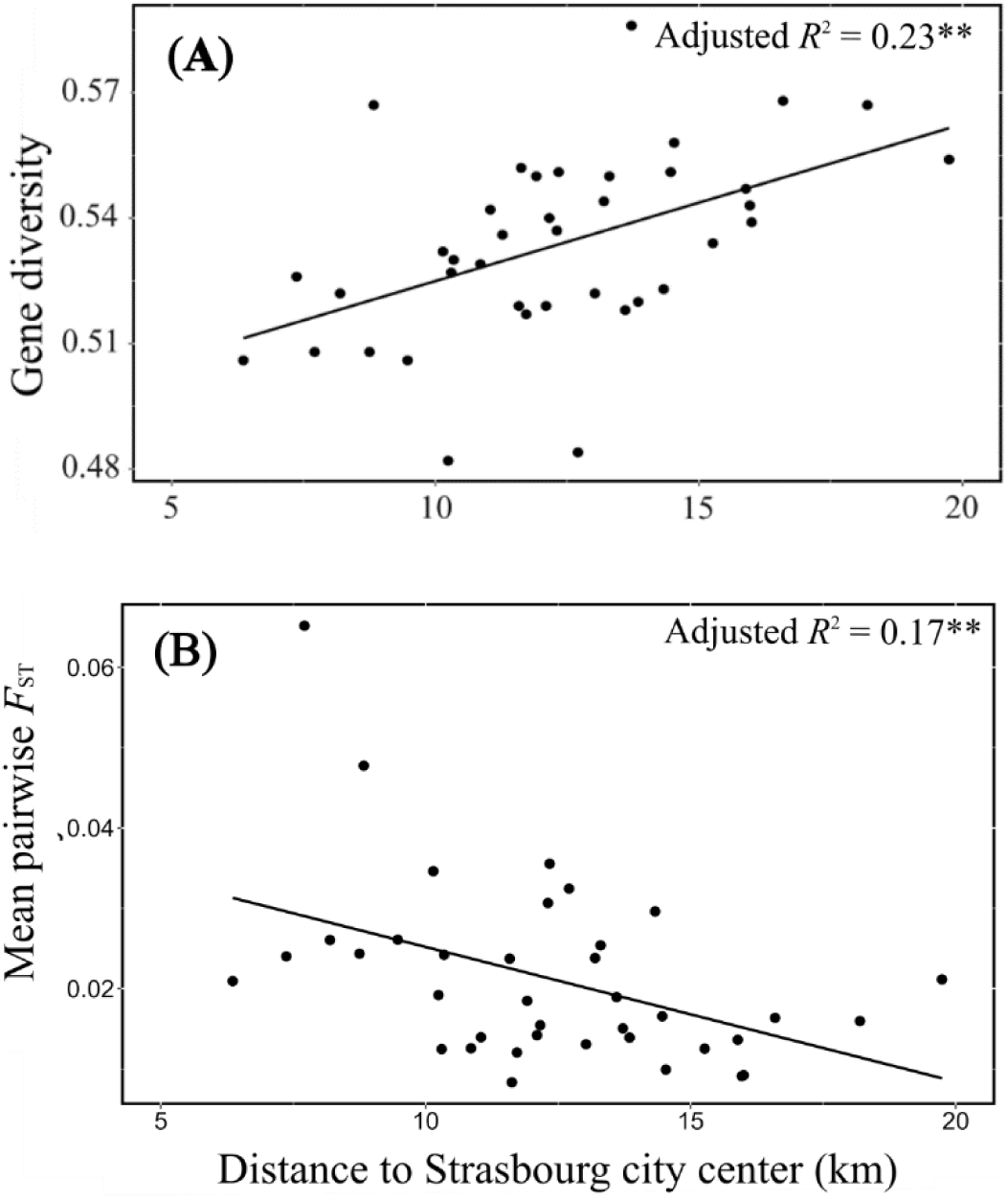
Relationship between the distance of populations to the centre of Strasbourg within a radius of 20 km buffer (see Figure S3) and the genetic diversity (*H*e) **(A)** and the mean pairwise *F*ST calculated according to Weir and Cockerham (1984) between each population and other populations within this radius **(B)**. Lines show the linear model. **: *P* < 0.01

Populations located in Northern France exhibited significantly lower levels of genetic diversity compared to those in Alsace, with a mean allelic richness of 2.91 ± 0.43, a mean *H*e of 0.43 ± 0.07, and a mean *H*o of 0.48 ± 0.07 (permutation tests, all at *P* <0.001; Table S1, Figure 2A, Figure S4). The mean intra-population kinship level in Northern France was 0.136 (Table S1, Figure 2B). Within Northern France, levels of genetic diversity and intra-population relatedness considerably varied among the different subregions (Figure S5 and Figure S7), with subregions N1, N2, and N6 showing levels similar to those observed in the Alsace subwatersheds, while the more isolated subregions N3 to N5 showed lower levels of genetic diversity (permutation tests, *P* < 10^-3^).

### Delimitation of population boundaries and genetic discontinuities

The sPCA showed contrasting geographic partitions of populations for the two studied areas: while a clear spatial structure appeared in Northern France, with genetic grouping corresponding to the major geographical regions, Alsacian populations were rather characterised by a gradient of population delimitation (Figure 4).

**Figure 4:**
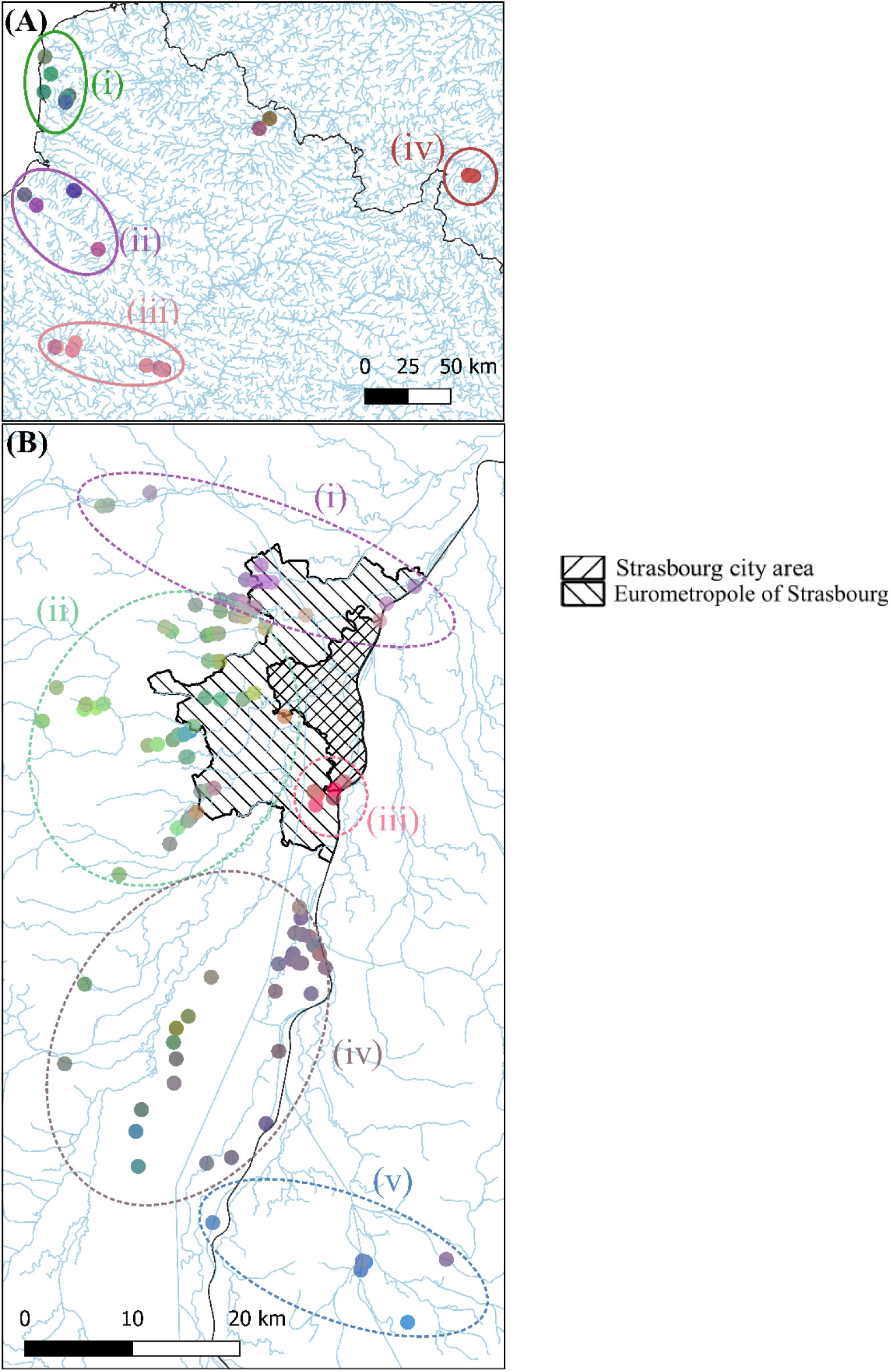
sPCA analyses depicting population genetic affiliation in Northern France **(A)** and Alsace **(B)**. Population colours reflect their coordinates on the first three axes of the sPCA, with coordinates along these three axes assigned to the Red, Green, and Blue channels respectively. Broad groupings of populations can be visualised by dashed labelled ellipses. Strasbourg city and the eurometropole of Strasbourg are represented by the striped areas. Main watercourses are also represented. Watercourses from COPERNICUS Land Monitoring Service, 2019: EU-Hydro.

In Northern France, the first three axes of the sPCA discriminated four distinct spatial groups: (i) populations located in subregion N3 in the north-western part, (ii) the populations located in subregion N5 in the south-western part, (iii) populations located in subregions N2 and N6 in the southern part of the sampling area, (iv) and finally populations located in subregion N1. The populations in subregion N4 showed a less clear assignment (Figure 4A).

In Alsace, five spatially structured groups could be distinguished by the first three axes of the sPCA (Figure 4B). These groups gather: (i) populations north of Strasbourg, mostly in the A6 subwatershed, (ii) populations west of Strasbourg, within subwatersheds A1, A2 and A5, (iii) populations to the north of subwatershed A4, just south of Strasbourg, (iv) populations of the remainder of subwatershed A4, further south from Strasbourg, (v) and populations located in subwatershed A3 and to the extreme south of subwatershed A4 (Figure 4B). As the boundaries between the different genetic groups did not show clear breaks, these groups reflected a gradient of genetic dissimilarity (Figure 4B).

### Levels of genetic differentiation and spatial genetic structure

Populations were highly genetically differentiated in northern France, with a mean multilocus *F*ST of 0.135 (95% confidence interval [0.113, 0.158]; Table S3, Figure S8A). Significant levels of population genetic differentiation were observed within most subregions (Table S3, Figure S8B). Accordingly, 91.9% of the 210 pairwise population *F*ST significantly differed from 0 after Bonferroni correction, most of the non-significant values of *F*ST being observed among populations belonging to the same geographical regions (Figure S9).

On the contrary, Alsacian populations were weakly, yet significantly genetically differentiated, with a mean multilocus *F*ST of 0.022 (95% confidence interval [0.020, 0.024]; Table S3, Figure S8A). Significant population differentiation was also observed within each subwatershed, except for very close populations in subwatershed A3 (Table S3, Figure S8B). Only 20.1% of the 3081 pairwise population *F*ST significantly differed from 0 after Bonferroni correction (Figure S8). Four populations (Gox1, Rlei45, S17, and S21, mapped on Figure S3) accounted for most of the significant pairwise *F*ST estimates (Figure S10).

When focusing on spatial patterns of genetic differentiation in Alsace, we observed a weak, yet statistically significant, pattern of isolation-by-distance (IBD), as shown by the positive relationship between pairwise population *F*ST/1-*F*ST and log-transformed Euclidean geographical distances (*rz* = 0.238, *P* < 0.01, Figure5A). However, this pattern goes along with an increase in pairwise *F*ST variance beyond a scale of ∼8 km, which suggested a predominant effect of drift beyond this scale of observation. Consistently, genetic relatedness among individuals decreased with increasing geographical distances, with significantly positive kinship coefficients occurring up to 11 km (Figure 5C). Interestingly, when distances among populations were calculated along watercourses (dStream), this pattern of IBD disappeared (*rz* = 0.045, *P* = 0.136, Figure 5B). However, positive *F*ij values still occurred for up to 20 km along streams (Figure 5D).

**Figure 5:**
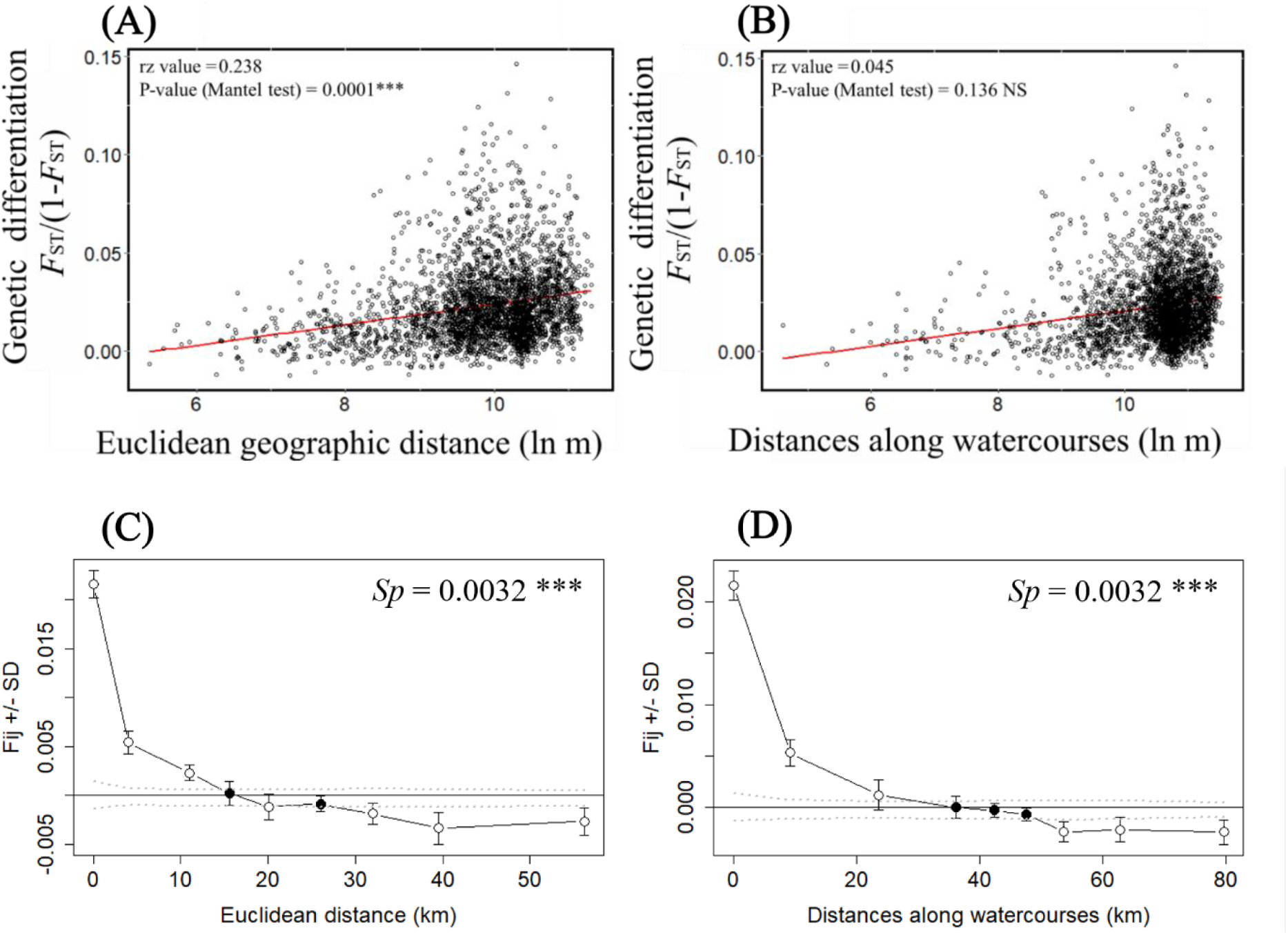
Spatial genetic structure in southern damselfly populations across the whole Alsace area. Relationship between levels of genetic differentiation (*F*ST/(1-*F*ST)) and log-transformed Euclidean distance (dEucli) **(A)** or watercourse geographical distance (dStream) **(B)**. Average pairwise kinship coefficients (*Fij*, Loiselle *et al*., 1995) between individuals are plotted against increasing classes of geographical Euclidean distances **(C)** or of waterways distance **(D)**. Standard errors for *F*ij for each distance class were obtained by jackknifing over loci. Dashed lines indicate the upper and lower 95% confidence intervals of non-significant spatial genetic structure, with white dots indicating significant estimates outside this confidence interval. *Sp* statistics based on the regression with ln(distance) and their statistical significance are also indicated.

Within subwatersheds, IBD was stronger using Euclidean distances compared to aquatic watercourse (dStream) distances in subwatersheds A1, A4 and A5 (respectively *rz* = 0.374, 0.414 and 0.472; *P* < 0.0001, *P* < 0.05 and *P* <0.0001; Figure S11). Within subwatersheds, there was a general short-distance spatial autocorrelation pattern, respectively up to 2.2 km, 4.6 km and <1 km in watersheds A1, A4 and A5 (Figure S11). A significant correlation between pairwise population genetic differentiation estimates and dStream was only found in subwatersheds A1 and A4 (Figure S11D-F). Yet, pairwise individual kinship coefficients decreased with aquatic distances (dStream), and were significantly positive for up to < 500 m, 4 km and about 700 m respectively in subwatersheds A1, A4 and A5 (Figure S12).

### Estimated Effective Migration Surface (EEMS)

The results with 1500 demes represented the best balance between precision and calculation time. The EEMS plot of effective migration rates identified an area of reduced migration rates covering most of the city of Strasbourg, and extending to the South (Figure 6). This notably encompasses populations located in the north of Strasbourg city (populations named S17, S21, CCR4, CCR2). Three additional areas showed lower migration rates than expected under an IBD at equilibrium. The first one was located downstream along subwatershed A5. The second was found in the midstream of subwatershed A1. The last area of restricted migration was centred on population Gox1, the population most upstream of subwatershed A2 (Figure 6).

**Figure 6:**
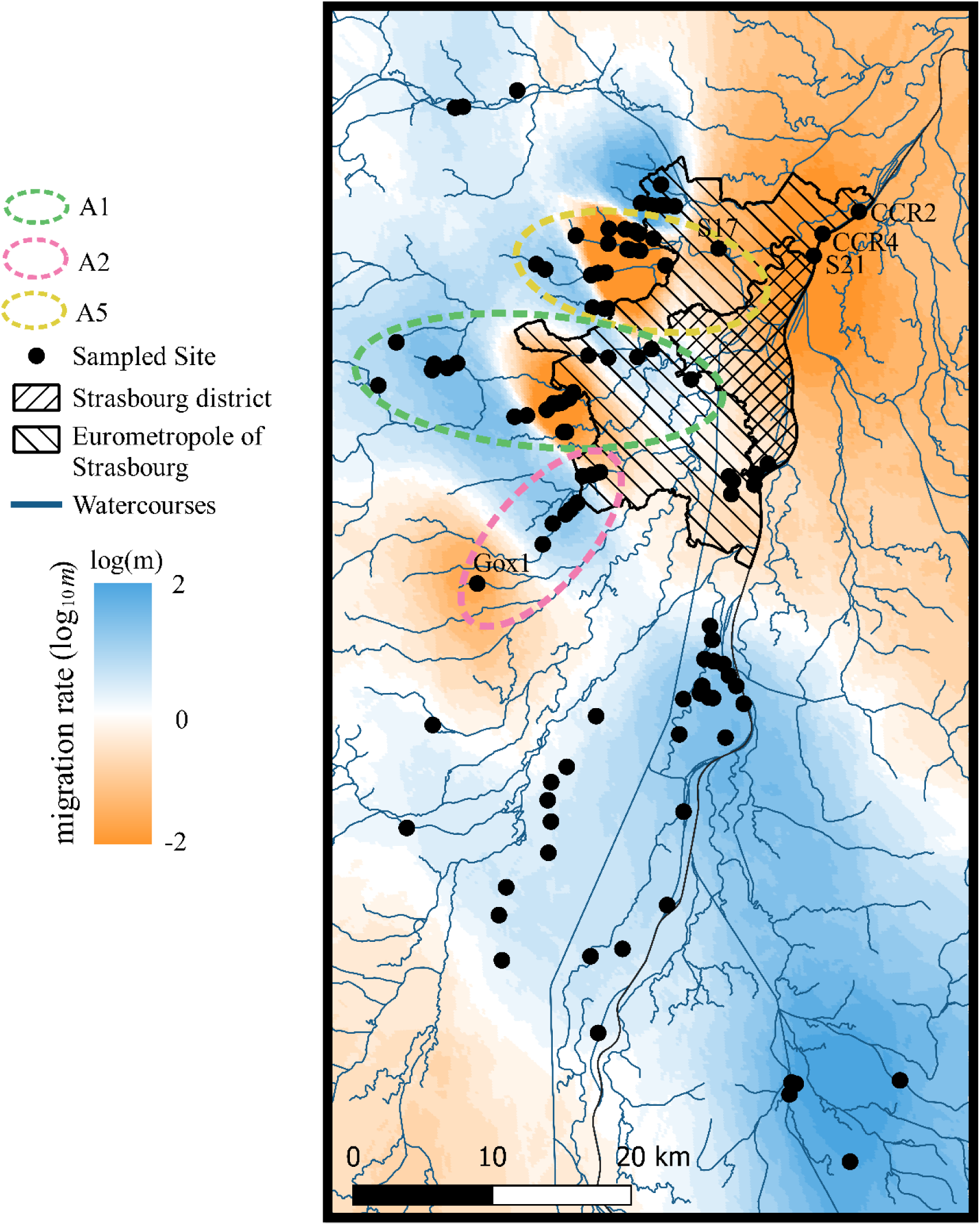
Estimated effective migration rates in southern damselfly in the Alsace region, as visualised by interpolated surface of the posterior mean migration rates *m* (on a log10 scale), where darker shades of blue indicate greater than expected migration (0-2), darker shades of orange indicate lower than expected migration (-2 to 0), and white indicates the null hypothesis of isolation-by-distance (0). The black circles indicate population locations. Main watercourses around Strasbourg City (blue lines) are also represented. The Strasbourg district area and the eurometropole of Strasbourg are represented by the striped areas. The locations and names of specific populations cited in the text (Gox1, S17, S21, CCR4, CCR2 were added to the map. Contours of the A1, A2, and A5 sub-catchments are indicated by dotted lines.

## Discussion

### Patterns of genetic diversity and population genetic differentiation between central and peripheral populations

Consistent with the first prediction of the “abundant centre” hypothesis (Eckert *et al*., 2008; Guo, 2012), peripheral populations in northern France showed significantly lower levels of genetic diversity as compared to central Alsacian populations. In addition, the two areas displayed different spatial distributions of genetic diversity: while indices of intra-population genetic diversity were constantly high over all populations located in Alsace, they widely differed between the surveyed subregions in Northern France. Indeed, at a finer scale in Northern France, populations within the most peripheral, isolated, and small habitat patches of the species distribution (N3, N4, N5) showed lower levels of genetic diversity than those from larger and more central patches (N1, N2, N6; Figure 1).

Also consistent with the second prediction of the “abundant centre” (Eckert *et al*., 2008; Pironon *et al*., 2017), the mean genetic differentiation in Alsace was lower than in Northern France. This pattern observed for marginal populations may be due to larger geographical scale of population separation in Northern France. Yet, similarly high levels of population differentiation were previously observed in other peripheral regions, even at small geographical scales (see Watts *et al*., 2004c, 2005; Lorenzo-Carballa *et al*., 2015). Indeed, indirect genetic approaches indicated dispersal up to 2 km and small-scale population genetic structuring (Watt *et al*., 2004c; 2006). Besides, the sPCA analysis could not reveal any major disjunction across the Alsace area. Both the low levels of population differentiation and low genetic relatedness within Alsacian populations support the idea that there is a substantial amount of gene flow among Alsacian populations (Ellegren and Galtier, 2016). By contrast, in Northern France, the sPCA analysis revealed striking genetic discontinuities between four large spatial groups of populations and suggested restricted gene flow among populations even at small scales, as exemplified by high levels of genetic differentiation observed within isolated sets of populations located in subregions N3, N4, N5.

In Northern France, gene flow may be reduced over short distances, as illustrated by population Bai3 in subregion N3, which showed low levels of genetic diversity and high levels of genetic differentiation with close neighbouring populations (< 5 km). Our findings thus reiterate the claim of Lorenzo-Carballa *et al*. (2015), namely that habitat management should improve connectivity with geographically close populations to prevent further loss of genetic diversity. Interestingly, no sign of inbreeding was detected, even in isolated populations or in populations characterised by low levels of genetic diversity, a result consistent with previous reports on southern damselflies (Watts *et al*., 2004c; Keller *et al*., 2012).

Altogether, the spatial distribution of genetic diversity and the patterns of genetic differentiation observed in the two regions are thus in agreement with the “abundant centre” hypothesis (Pironon *et al*., 2017). This pattern, observed towards the northern limit, may exist in other cardinal directions as well, because peripheral populations of the southern damselfly are declining in their northern, eastern (Grand, 1996) and southernmost limits in North Africa (Ferreira *et al*., 2015). It may also result from colonisation history (see Johansson *et al*., 2013), and more extensive sampling would certainly provide a more global view of the evolution of genetic structuring in populations of the southern damselfly.

### Spatial genetic structure and dispersal pathways

Although the level of genetic differentiation among southern damselfly populations was weak across the Alsace area, we found an isolation-by-distance (IBD) pattern. This pattern was reflected both by an increase in genetic differentiation among populations and a decrease of pairwise individual kinship with increasing Euclidean geographical distances. This result is consistent with previous studies that observed an IBD pattern in southern damselfly populations both at a fine scale (Watts *et al*., 2004c; Watts and Thompson, 2012; Lorenzo-Carballa *et al*., 2015) and at a broader scale of investigation (Watts *et al*., 2006; Watts and Thompson, 2012).

Population genetic structure also depends on the biological attributes of a species, in particular its dispersal capabilities and habitat requirements (Hutchison and Templeton, 1999; Phillipsen *et al*., 2015; Ellegren and Galtier, 2016). The relationship between genetic and geographic distances across the Alsace area suggested that gene flow events among southern damselfly populations may occur in a stepping-stone pattern among patches of suitable habitats (Kimura, 1953; Kimura and Weiss, 1964; Hutchison and Templeton, 1999), consistent with the patchy distribution of habitats of this species (Conrad *et al*., 1999; Rouquette and Thompson, 2005; Purse and Thompson, 2009). Adults can disperse among suitable habitat patches, but since they are poor fliers (Watts *et al*., 2004c), dispersal is likely to occur mostly at short distances between neighbouring populations.

The IBD patterns were stronger at the smaller scale of subwatersheds compared to the one observed across the whole Alsace region. In addition, variance in pairwise genetic differentiation in Alsace strongly increased beyond geographical distances larger than 8 km. These results suggest that at large geographical distances, gene flow among populations are influenced by ecological factors additional to the simple Euclidean distances, suggesting potential barriers to gene flow occurring over large scales over 8 km (e.g. forested or elevated areas; Hutchison & Templeton, 1999; Phillipsen *et al*., 2015).

In this respect, the geographical distances strictly calculated along watercourses only partly explained the levels of genetic differentiation observed among populations. This suggests that dispersal events can occur overland and is not restricted to stream networks. This overland dispersal between different streams is further supported by the sPCA analysis that suggested no clear genetic discontinuity between the different river networks. These results are in line with the results of Keller and Holderegger (2013), where different movement strategies were described in southern damselflies for short- and long-distance dispersal: dispersal along watercourses appeared to be favoured over short distances (< 3 km), whereas dispersal along straight lines was best supported for long distances (> 3 km). Altogether, a simple IBD is not likely to explain the totality of the spatial genetic structure we observed in Alsace, pinpointing the possible role of landscape features in shaping dispersal among populations (Storfer *et al*., 2007).

### Urbanisation effects on spatial genetic structure

The city of Strasbourg likely acts as a barrier to gene flow between southern damselfly populations in Alsace. Indeed, the EEMS analysis clearly showed a reduced migration rate in the vicinity of the city. This result was further supported by increasing levels of population genetic differentiation and intra-population kinship coefficients in the vicinity of the city of Strasbourg. A typical “urban fragmentation” pattern may thus be observed; urbanised areas appeared to fragment habitats, reducing gene flow and increasing genetic differentiation among populations (Lorenzo-Carballa *et al*., 2015; Johnson and Munshi-South, 2017; Lourenço *et al*., 2017; Miles *et al*., 2019). However, while the city of Strasbourg represented a clearly restricted migration zone, peripheral small towns and villages did not suggest any major barriers to gene flow. Therefore, the impact of urbanisation on patterns of gene flow likely differs depending on the size and level of urbanisation (e.g. Trumbo *et al*., 2019). The fragmentation impact of the city of Strasbourg can extend to a large distance outside the city centre, because the urban area extends over more than 300 km², up to 12 km away from the city centre. Further away from the city, restricted migration among populations may be due to other landscape ecology features, such as forest patches or high-elevation areas. As a matter of fact, the significant relationship observed between levels of genetic diversity and genetic differentiation and the geographical distance from the city disappeared beyond a scale of 20 km. This finding goes along with a general increase in pairwise *F*ST variance with increasing geographical distance, which suggested additional landscape effects other than urban areas, such as topography and land-use (Hutchison and Templeton, 1999; Storfer *et al*., 2007).

Moreover, the “urban fragmentation” model further postulates that fragmentation and isolation of populations by unfavourable urbanised areas could increase the strength of genetic drift, which erodes neutral genetic diversity within populations (Miles *et al*., 2019). Levels of allelic richness and gene diversity indeed decreased in the direct neighbourhood of Strasbourg city, thus suggesting a negative relationship between urbanisation and genetic diversity, as it was described in several species (Noël *et al*., 2007; Munshi-South *et al*., 2016; Johnson and Munshi-South, 2017; Miles *et al*., 2019; Kimmig *et al*., 2020). Nonetheless, a reduction in gene flow linked to urbanisation is not systematically associated with a reduction in genetic diversity (Trumbo *et al*., 2019; Fusco *et al*., 2021): indeed, population S21 surprisingly showed high levels of genetic diversity despite a substantial genetic differentiation with close populations and its proximity to Strasbourg city. This could be explained by incoming migrant individuals from diverse sources outside our sampling area (Whitlock and McCauley, 1990).

Our findings on city impacts on population genetic features again require caution before being generalised, and would need to be repeated in different urban areas in order to confirm that populations evolve in the same way in response to urbanisation (see Santangelo *et al*., 2018; Miles *et al*., 2019; Diamond and Martin, 2021). Incorporating landscape components among populations, along with the use of more resolutive genome-wide molecular markers, should be pursued to better understand the impact of urbanisation and landscape features on southern damselfly populations at a fine scale. This may also help to explain the occurrence of zones of restricted migration observed in rural areas. Furthermore, cities can exhibit considerable spatial and temporal heterogeneity at fine scale, which could influence population adaptation (Rivkin *et al*., 2019). It would therefore be useful to study the adaptive variation that can influence the evolution of species in urban environments (Johnson and Munshi-South, 2017; Rivkin *et al*., 2019; Diamond and Martin, 2021; Babik *et al*., 2023). Indeed, in another species of damselfly, *Coenagrion puella,* urbanisation and associated heat islands seem to select for slower development and higher survival (Tüzün *et al*., 2017). Likewise, in southern damselflies, development time is also affected by water temperature associated with the release of industrial cooling waters (Thelen, 1992).

### Conclusion

Central populations of southern damselflies located in urbanised environments in Alsace exhibited higher levels of genetic diversity and lower levels of genetic differentiation compared to northern peripheral populations, indicating that the species might follow a classical “abundant centre” model in terms of population genetic structure (Guo, 2012; Pironon *et al*., 2017). Populations found in isolated suitable habitats located at high latitudes exhibited particularly low levels of genetic diversity, which is alarming with regard to their evolutionary potential and may threaten their long-term persistence. In contrast, central Alsacian populations displayed a metapopulation structure with high intra-population levels of genetic diversity and a high amount of gene flow among populations. These findings contrast with previous studies on southern damselflies that suggested very low dispersal capabilities and high levels of genetic differentiation at fine scales of investigation (Watts *et al*., 2004c; Watts and Thompson, 2012; Lorenzo-Carballa *et al*., 2015). The highly urbanised city of Strasbourg likely represents a substantial barrier to gene flow among southern damselfly populations. Although this barrier effect of urban areas on southern damselfly dispersal has already been suggested by previous work conducted at a smaller scale (Lorenzo-Carballa *et al*., 2015; Watts *et al*., 2004c), our study is the first to clearly show a progressive decrease in genetic diversity in populations near a major urban area. The impact of urbanisation may also vary according to the type and size of urban areas encountered, as smaller Alsacian villages did not seem to impede gene flow among populations. Finally, Alsacian populations matched a classical pattern of isolation-by-distance population genetic structure. This suggests that individuals do not disperse exclusively along watercourses, their specific breeding habitat, but are also able to disperse over land. However, Euclidean distance alone did not fully explain the levels of genetic differentiation and the impact of the different landscape features occurring in the Alsace region remains to be determined. A genomic approach using SNP markers coupled with a landscape analysis would provide a more detailed and comprehensive vision of the evolutionary processes influencing patterns of gene flow and levels of genetic diversity occurring in this area of the southern damselfly geographical distribution.

## Acknowledgements

We thank SOCOS for financial support. We also thank the members of the CEN involved in the sampling of southern damselfly legs in the Hauts-de-France and Belgium. We also wish to express our gratitude to Anaëlle Meunier, Hana Mohoric, Clément Gardin and Louise Laine for help with sampling in Alsace. We are grateful to Xavier Vekemans, Eric Petit, and Frédéric Austerlitz for helpful discussions and suggestions related to this project. We thank Sandrine Belingheri for administrative support and Laurence Debacker for technical support.

## Biosketch

This work is a part of the PhD project of Agathe Lévêque which focuses on population genetics in the southern damselfly (*Coenagrion mercuriale*). She is interested in studying the spatial genetic structure and patterns of gene flow in the context of the anthropogenic impacts of urban areas and newbuild bypass highways. Her research approach is rooted in the field of conservation biology. All authors have a diverse range of expertise in population genetics, naturalist expertise and conservation biology.

## Author contributions

A.D, V.V, J-F.A conceived and designed the study; A.L, A.D, V.V, J-F.A and C.V organised the collection of specimens; A.L, A.D, F.D, C.V and J-F.A collected the samples in the field; A.L, F.D and C.G performed the molecular laboratory works; A.L. analysed the data; A.L, A.D and J-F.A wrote the first draft of the manuscript. All authors contributed to the final version of the manuscript and approved the submitted version.

## Data Storage

The dataset used, southern damselflies microsatellite genotypes and their sampling location will be deposited on Dryad upon acceptance of the article.

## Ethics approval statement

Capture permits were issued by DREAL du Grand Est and DREAL des Hauts-de-France after approval from the CSRPN Grand Est (Arrêté N°2021-DREAL-EBP-0104) and the CSRPN Hauts-de-France (Arrêté N°2021-ESP-24).

## Supporting information

Article entitled Contrasting patterns of spatial genetic structure in endangered southern damselfly (*Coenagrion mercuriale*) populations facing habitat fragmentation and urbanisation.

**Figure S1:**
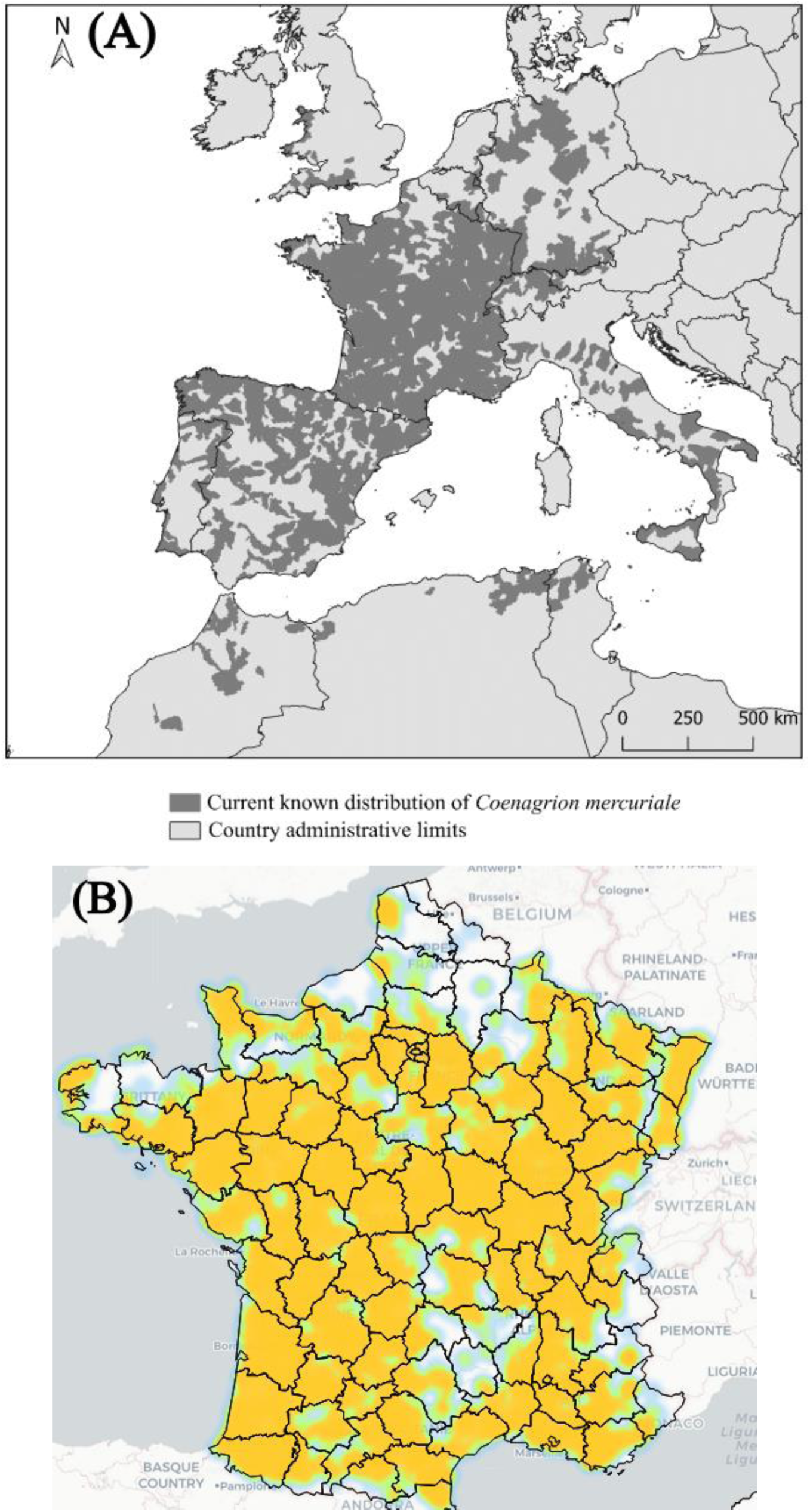
**(A)** Map showing the current known distribution (dark grey zones) of the southern damselfly (*Coenagrion mercuriale*). Data obtained from IUCN SSC Odonata Specialist Group 2019. The IUCN Red List of Threatened Species. Version 2022-2. https://www.iucnredlist.org/ Downloaded on 28 July 2023. Black lines indicate the administrative limits of countries. **(B)** Interpolation of the density of observations of southern damselflies in France. Source: https://atlas-odonates.insectes.org/odonates-de-france/coenagrion-mercuriale.

**Figure S2:**
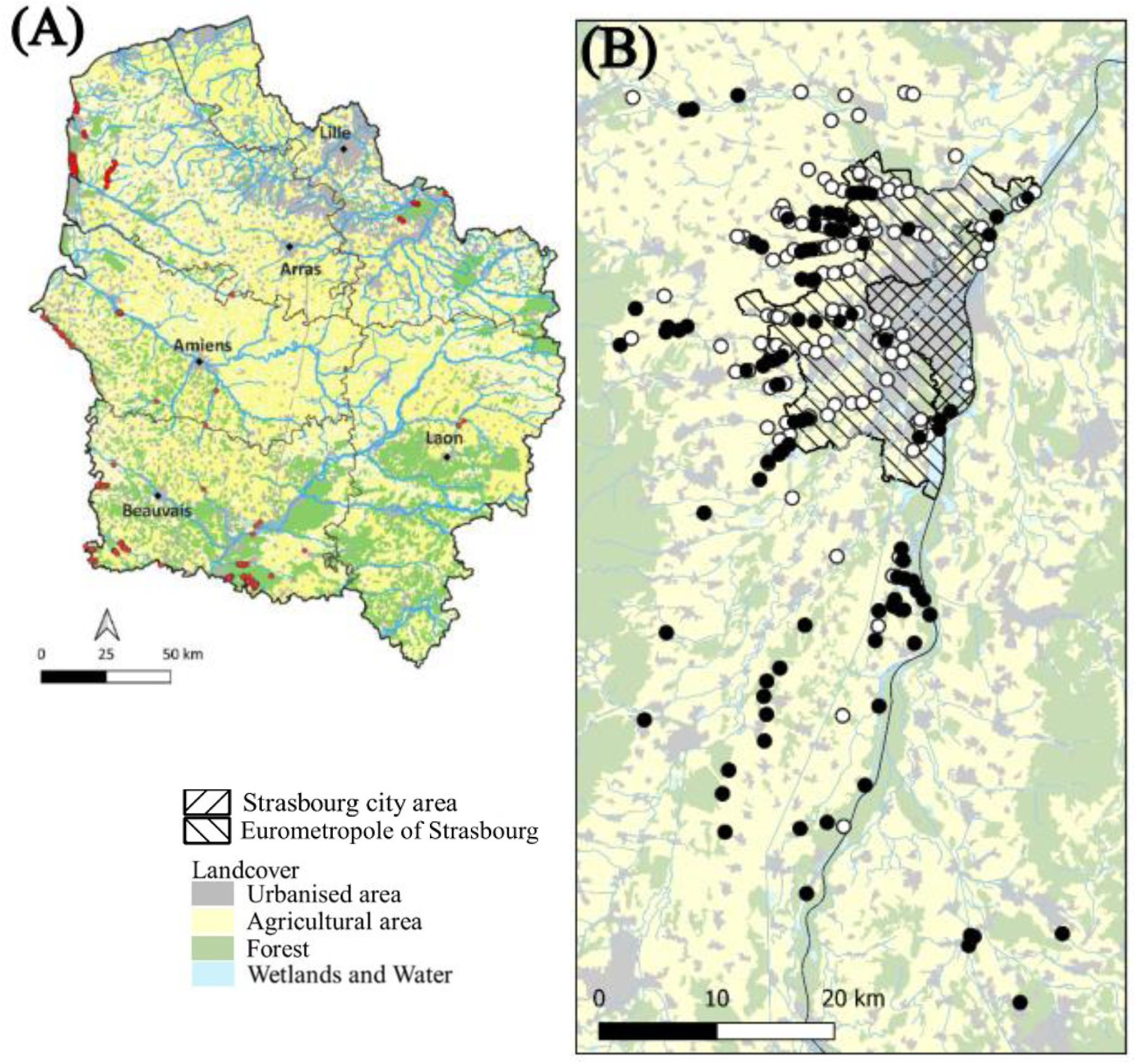
**(A)** Map showing the geographic distribution of southern damselfly in the region Hauts de France. The sites where the species is present appear in red. Blue lines; rivers; light blue regions; wetlands; yellow; agricultural zones; green: forest; grey: urban areas (https://irpn.drealnpdc.fr/wp-content/uploads/2023/04/PRA_20230420_DOC_PRA_Libellules_HdF.pdf.). **(B)** Geographical location of southern damselfly sampling sites around Strasbourg City from 2021 to 2022. Black dots represent sites successfully sampled, and white dots represent sites surveyed but where no southern damselflies were found. Main watercourses were taken from COPERNICUS Land Monitoring Service (2019: EU-Hydro). Strasbourg city and the eurometropole of Strasbourg are represented by the striped areas. Land cover was simplified from Corine Land Cover Edition 2018. Note the difference in spatial scales among the regions.

**Figure S3:**
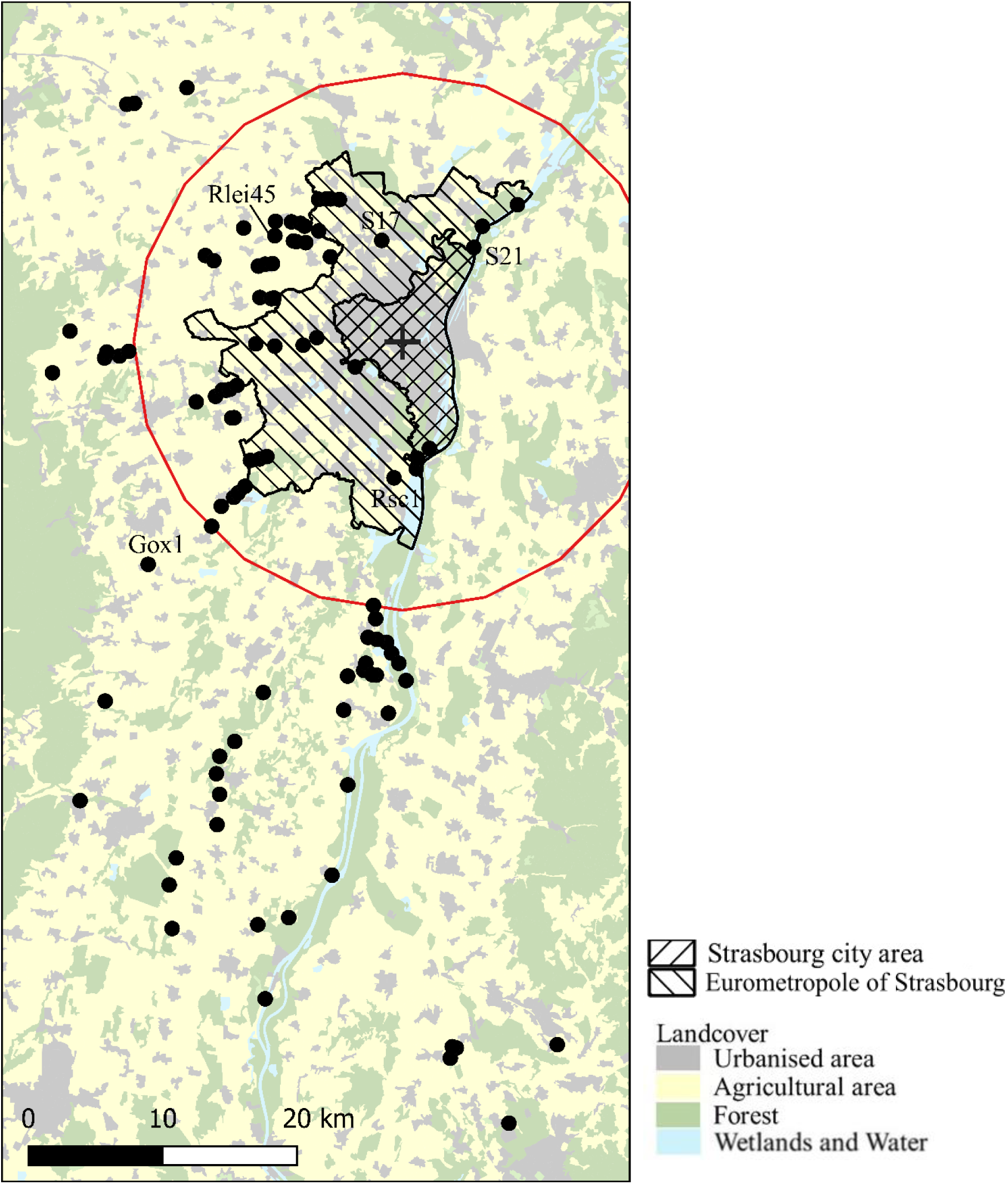
Map showing the 20 km radius (red line) around Strasbourg centre (black cross) within which the potential effect of the city of Strasbourg on southern damselfly populations was measured in terms of levels of genetic diversity (*A*r, *H*e), the mean levels of individual kinship coefficient (*F*ij) and the mean pairwise levels of *F*ST calculated according to Weir and Cockerham (1984). Black dots represent sites sampled in Alsace with at least eight individuals. Land cover was simplified from Corine Land Cover Edition 2018 France Métropolitaine. Locations and names of specific populations cited in the text (Gox1, S17, S21, Rlei45, and Rsc1) were added to the map. Strasbourg city and the eurometropole of Strasbourg are represented by the striped areas.

**Figure S4:**
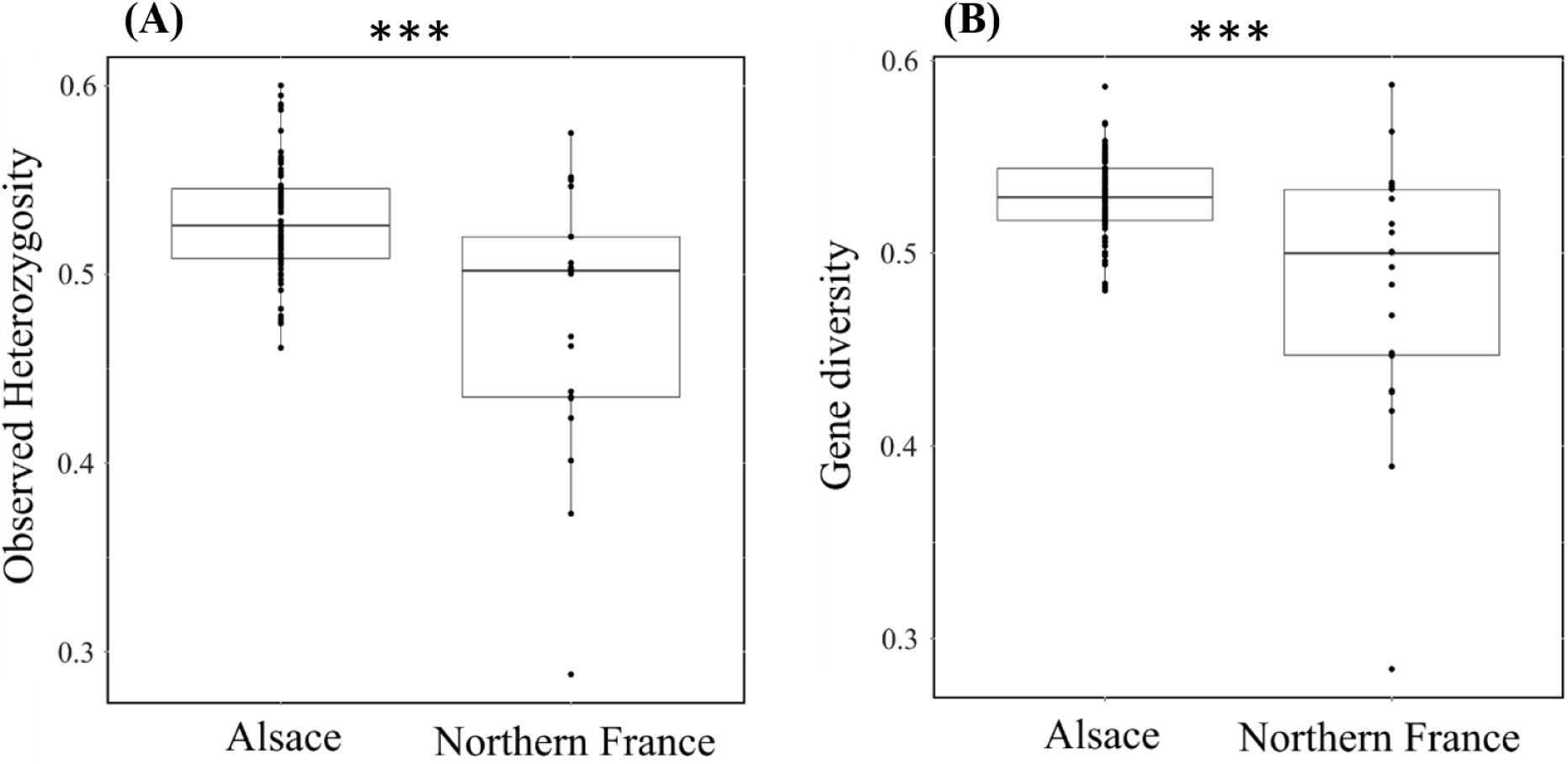
Distribution of observed heterozygosity *H*o **(A)** and gene diversity *H*e **(B)** within Alsace and northern France areas. *P*-value: *** *P* < 0.001.

**Figure S5:**
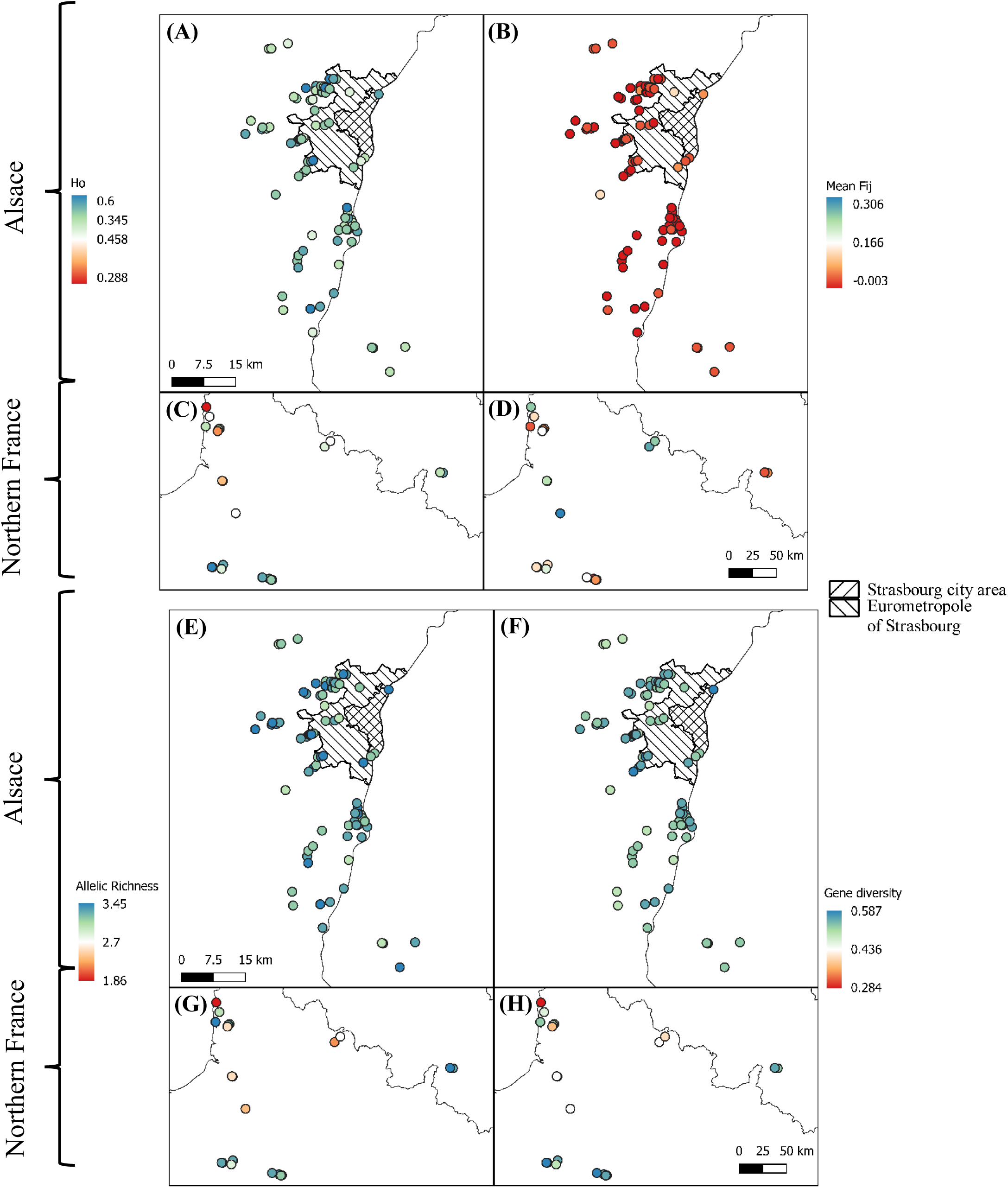
Geographical distribution of levels of observed heterozygosity (*H*o) **(A, C)**, levels of intra-population kinship coefficient *F*ij **(B, D)**, levels of allelic richness (*A*r) **(E, G)**, and levels of expected heterozygosity (*H*e) **(F, H)** in southern damselfly populations genotyped for at least eight individuals in Alsace **(A, B, E, F)** and Northern France **(C, D, G, H)**. Strasbourg city and the eurometropole of Strasbourg are represented by the striped areas.

**Figure S6:**
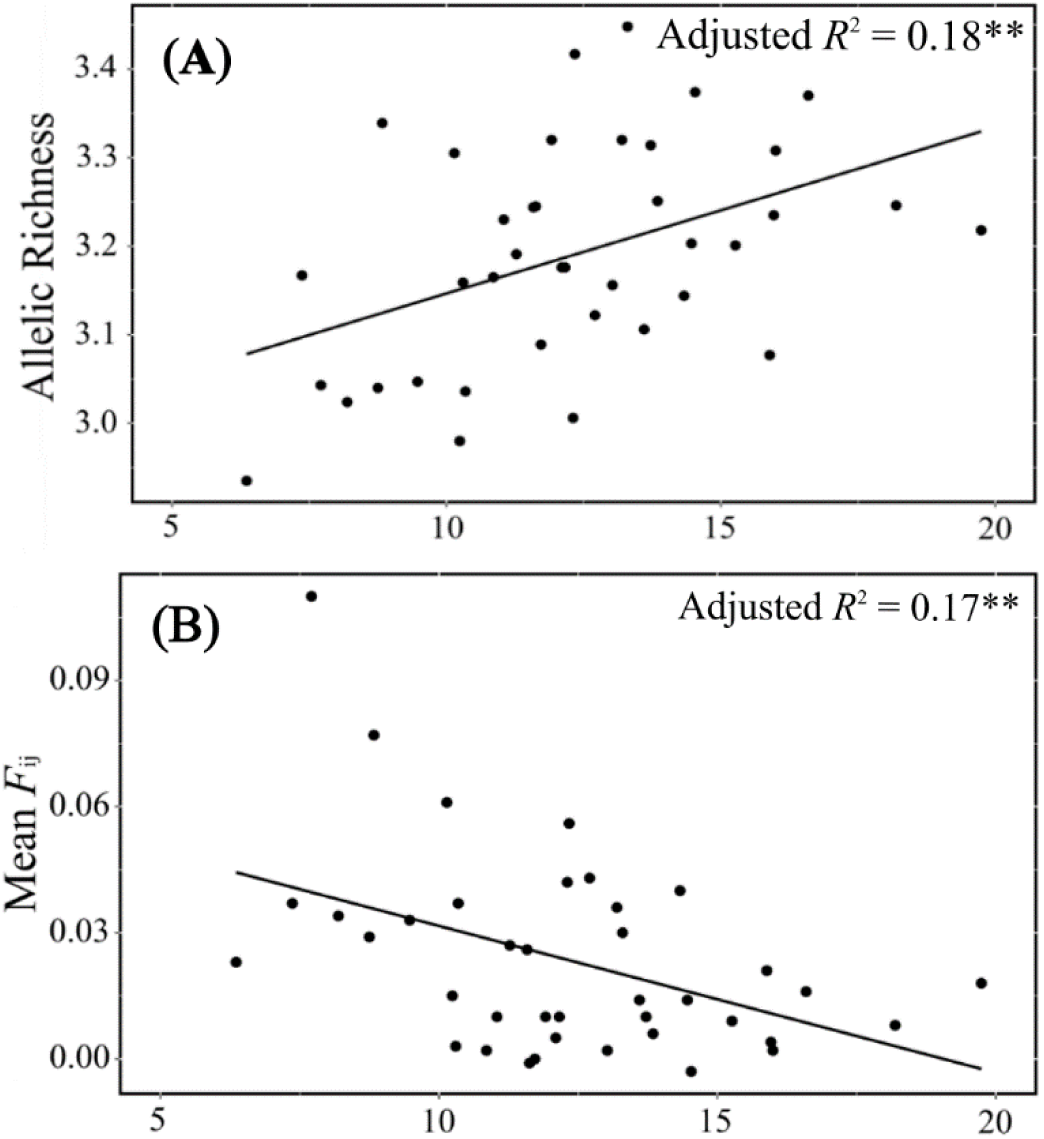
Relationship between allelic richness (*Ar*) **(A)** and mean intra-population kinship coefficient (*F*ij) **(B)**, and the distance of the population to the centre of Strasbourg within a 20 km buffer (see Figure S3). Lines show the linear model. *\*\*: P < 0.01*

**Figure S7:**
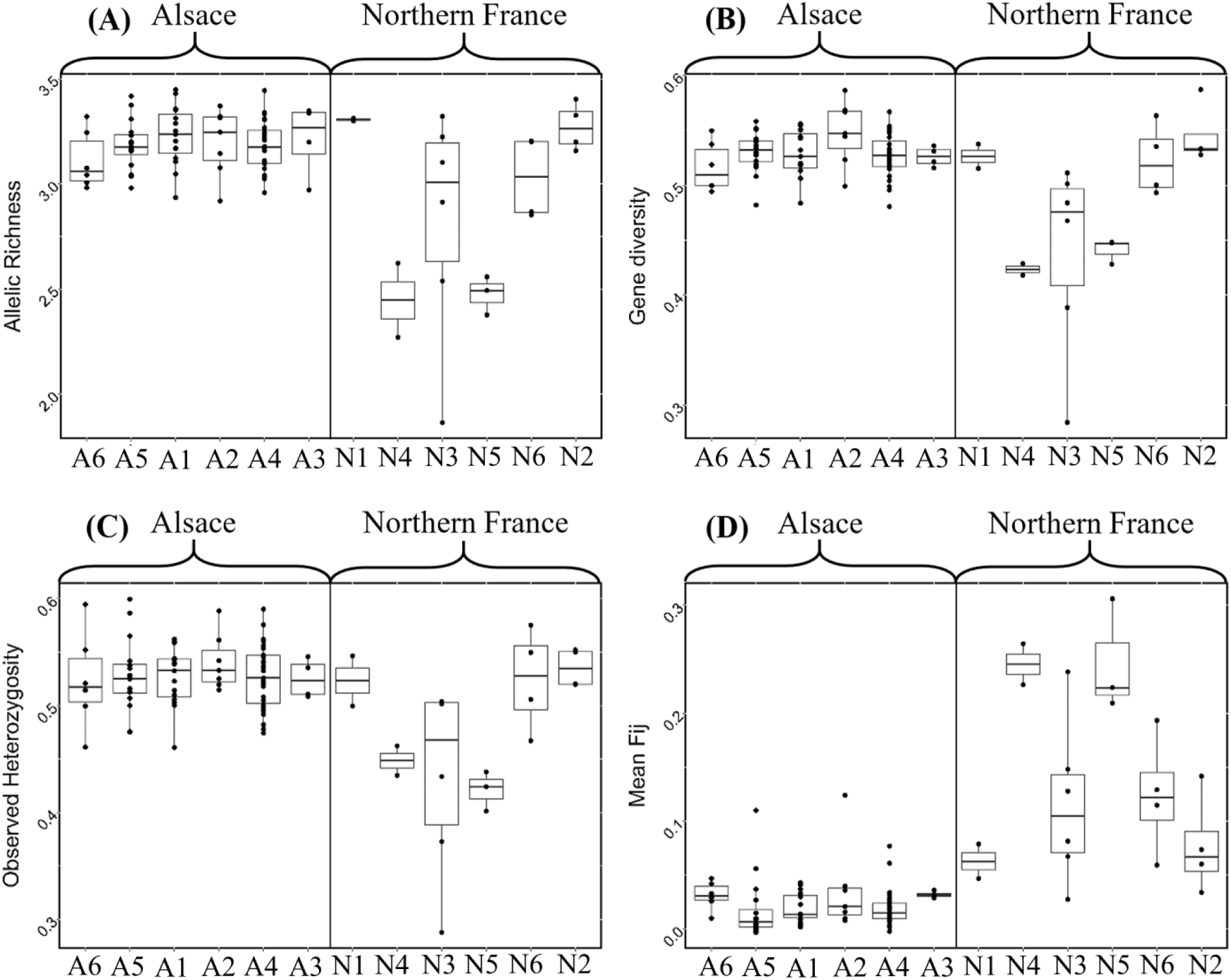
Distribution of allelic richness *A*r **(A)**, expected heterozygosity *H*e **(B)**, observed heterozygosity *H*o **(C)**, and intra-population kinship coefficient *F*ij **(D)** within each hydrographic unit or spatial regions, for populations with at least eight sampled individuals.

**Figure S8:**
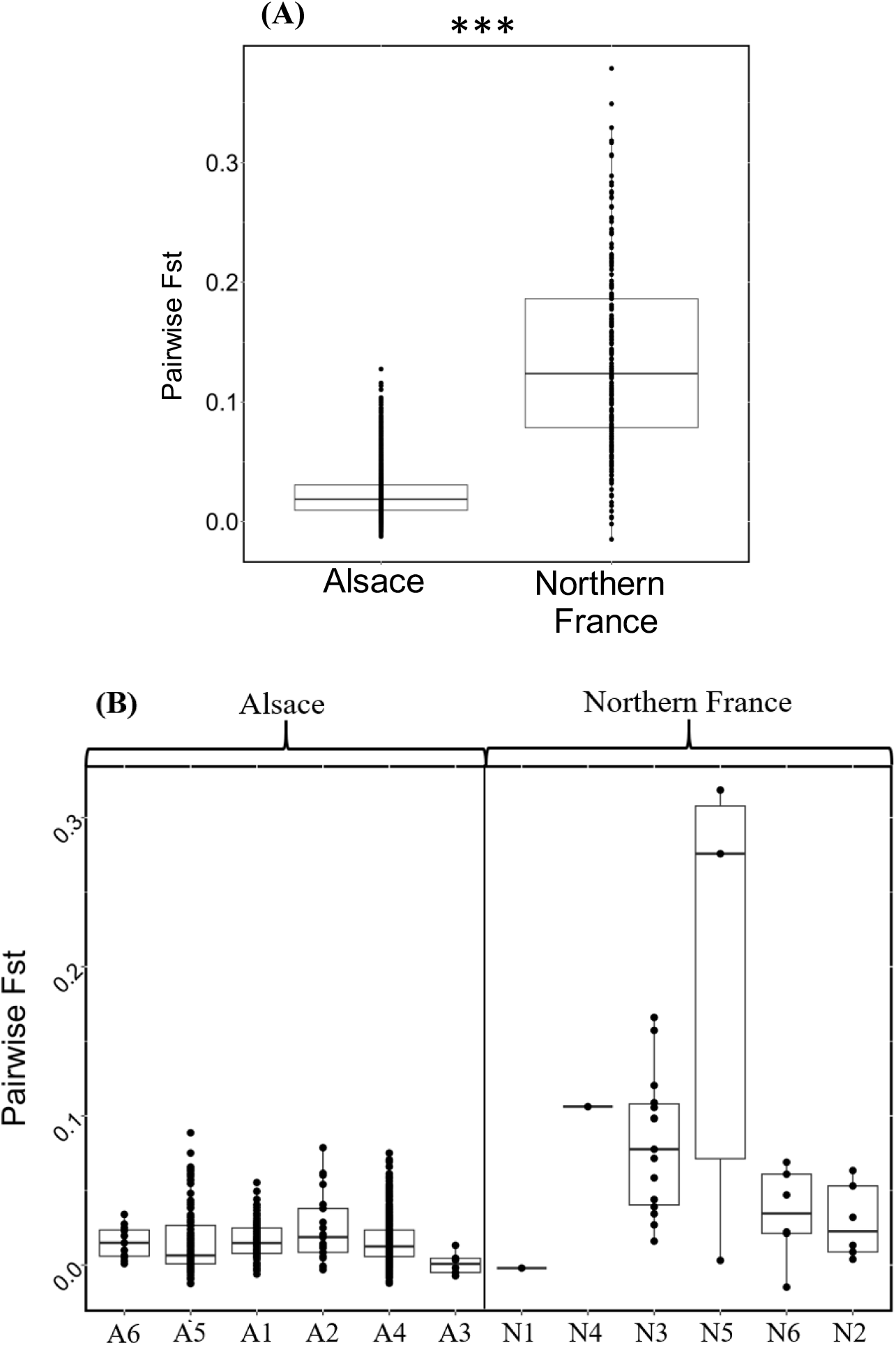
Distribution of pairwise *F*ST estimates within the two main surveyed areas, Alsace and Northern France **(A)** and within hydrographic units or spatial regions within both areas **(B)**, for populations with eight or more sampled individuals. *P*-value: *** *P* < 0.001.

**Figure S9:**
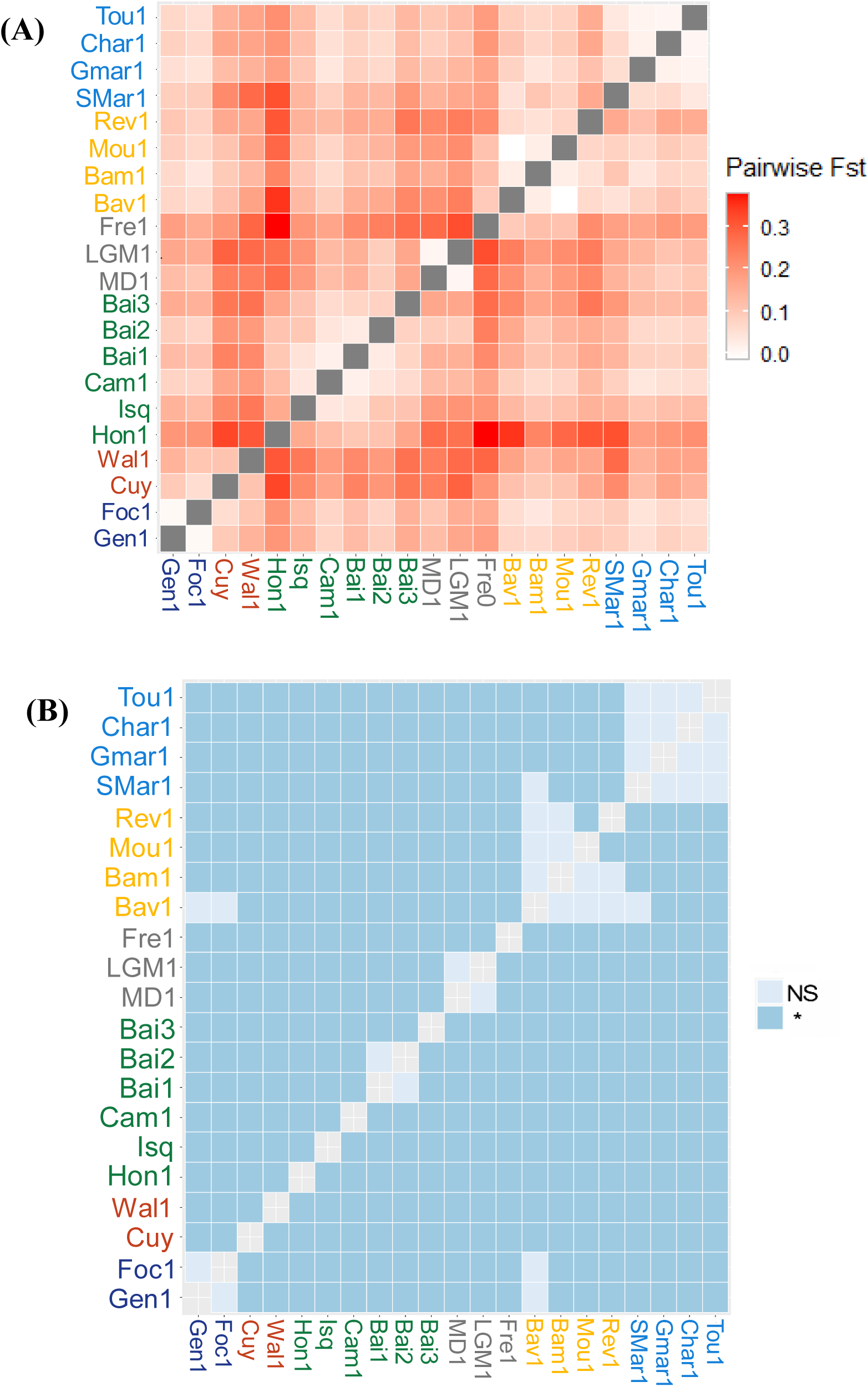
Matrices of pairwise *F*ST estimates in Northern France **(A)** and their associated statistical significance assessed using 10000 permutations of multilocus genotypes among populations after Bonferroni correction **(B)**. Populations are listed and coloured by geographical regions as in Figure 1.

**Figure S10:**
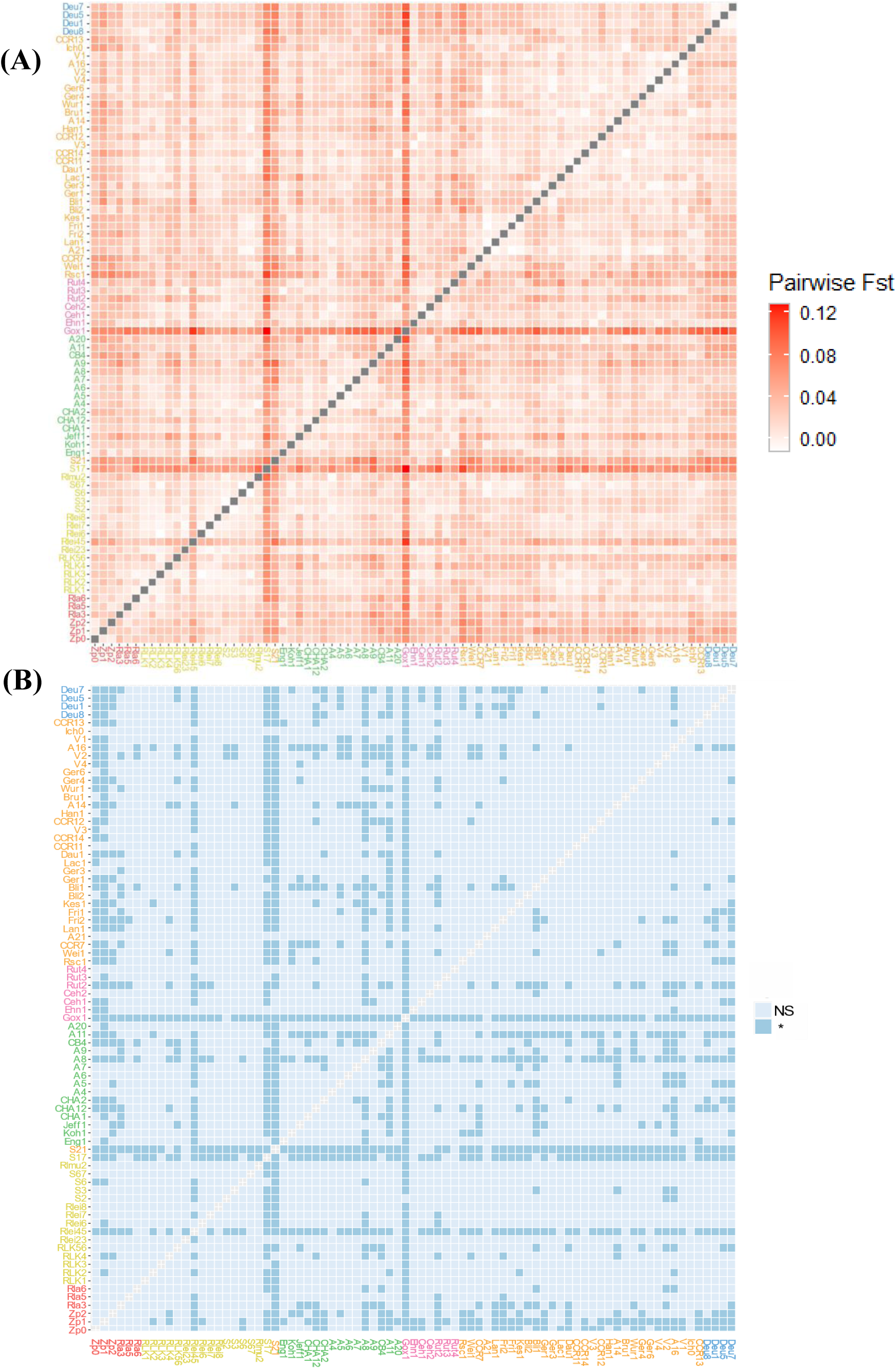
Matrices of pairwise *F*ST estimates in Alsace **(A)** and their associated statistical significance assessed using 10000 permutations of multilocus genotypes among populations after Bonferroni correction (**B**). Populations are listed and coloured by subwatershed as in Figure 1.

**Figure S11:**
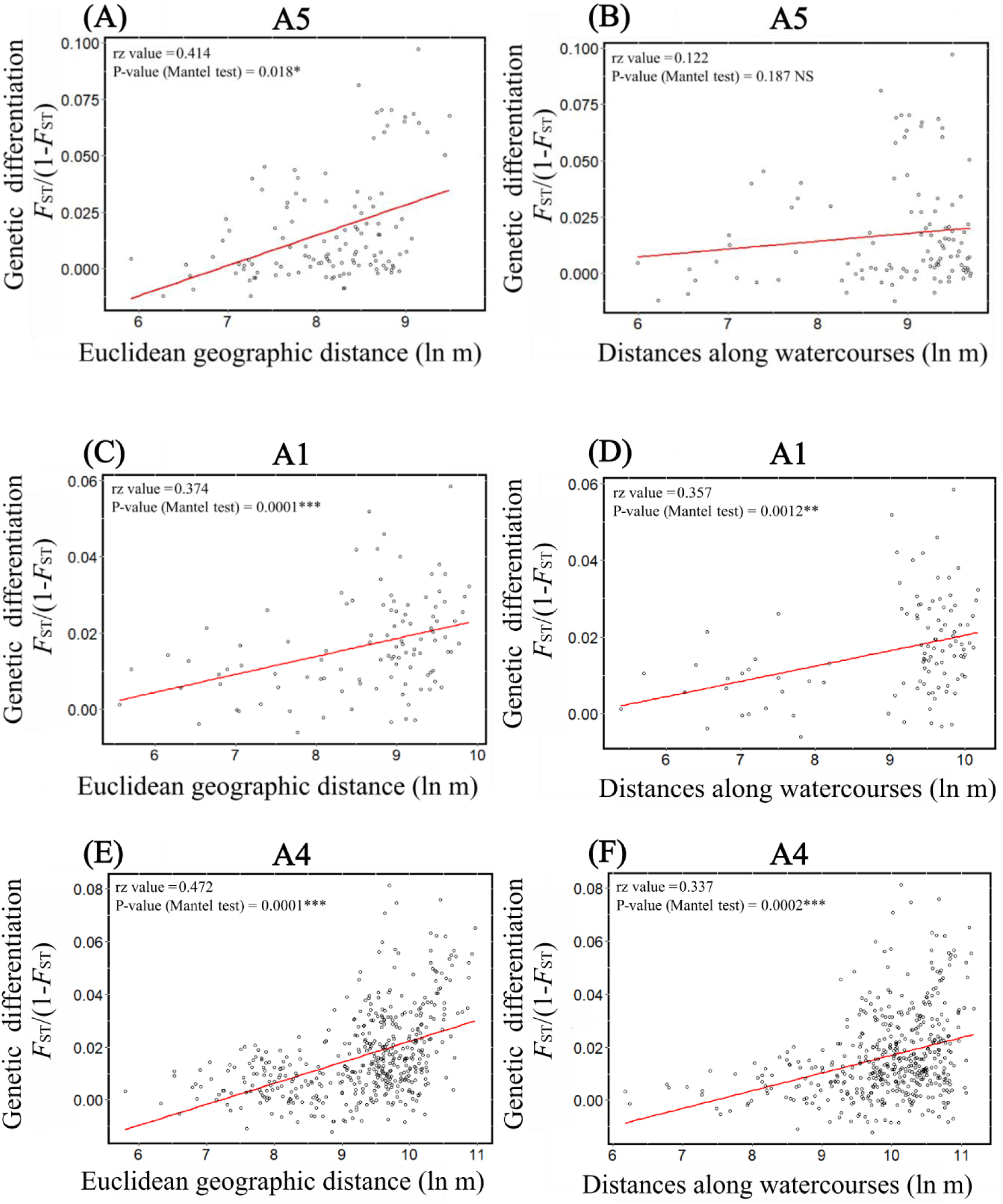
Relationship between levels of genetic differentiation (*F*ST/(1-*F*ST)) and log-transformed Euclidean distance (dEucli) **(A, C, E)** or watercourse geographical distance (dStream) **(B, D, F)** among populations of *Coenagrion mercuriale* within three main subwatersheds: A5 **(A, B)**, A1 **(C, D)**, A4 **(E, F)**.

**Figure S12:**
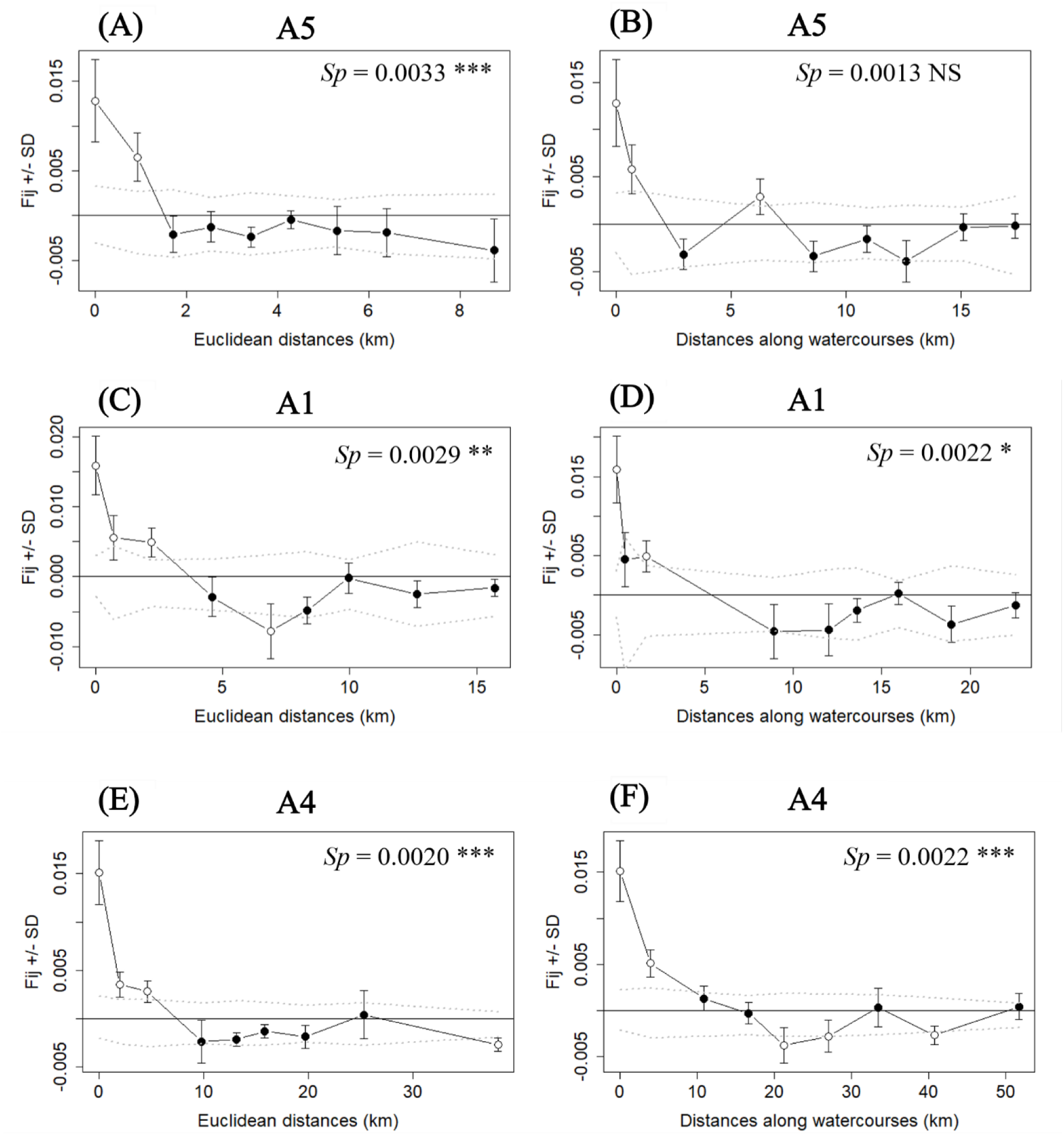
Broad spatial genetic structure in southern damselfly populations in three different main subwatersheds: A5 **(A, B)**, A1 **(C, D)**, A4 **(E, F)**. Average pairwise kinship coefficients (*Fij*, Loiselle *et al*., 1995) between individuals are plotted against increasing classes of geographical Euclidean distances **(A, C, E)** or of waterways distance **(B, D, F)**. Standard errors for *F*ij for each distance class were obtained by jackknifing over loci. Dashed lines indicate the upper and lower 95% confidence intervals of non-significant spatial genetic structure, with white dots indicating significant estimates outside this confidence interval. *Sp* statistics based on the regression with ln(distance) and their statistical significance are also indicated, allowing for comparisons of the strength of spatial genetic structure.

**Table S1:** Sampling locations and basic genetic diversity estimates for each studied population. ID: population name, Sampling Year: the year of sampling, Area: sampling region and hydrographic units they belong to, geographic coordinates of the sampling site (WGS84, EPSG4326), N: sample size. Summary of population structure for nuclear polymorphism at 10 microsatellite loci for each population can also be found: the total number of alleles (*A*T), the allelic richness (*A*r) rarefied on eight individuals, the observed heterozygosity (*H*o), the expected heterozygosity (*H*e), the intrapopulation fixation index *F*IS, and the mean intrapopulation kinship coefficient *F*ij. The significance of *F*IS was tested with 10000 permutations of alleles among individuals within populations (*: *P* < 0.05, **: *P* < 0.01, ***: *P* < 0.001), note that none was significant. -: no estimates because of insufficient population size.

| ID | Sampling Year | Area | Hydrographic units/ Spatial Region | Longitude | Latitude | N | $A_T$ | $A_r$ | $H_o$ | $H_e$ | $F_{IS}$ | $F_{ij}$ |
| --- | --- | --- | --- | --- | --- | --- | --- | --- | --- | --- | --- | --- |
| A11 | 2022 | Alsace | A1 | 7.655745 | 48.574643 | 39 | 41 | 3.167 | 0.533 | 0.526 | -0.013 | 0.037 |
| A14 | 2021 | Alsace | A4 | 7.697191 | 48.354712 | 27 | 39 | 3.073 | 0.526 | 0.528 | 0.004 | 0.008 |
| A16 | 2022 | Alsace | A4 | 7.653206 | 48.218936 | 34 | 40 | 3.178 | 0.547 | 0.552 | 0.009 | 0.026 |
| A20 | 2022 | Alsace | A1 | 7.669929 | 48.579161 | 13 | 32 | 2.935 | 0.523 | 0.506 | -0.034 | 0.023 |
| A21 | 2022 | Alsace | A4 | 7.595143 | 48.343877 | 9 | 31 | 3.029 | 0.478 | 0.481 | 0.006 | 0.011 |
| A3 | 2021 | Alsace | A1 | 7.532710 | 48.540547 | 1 | 17 | - | - | - | - | - |
| A4 | 2022 | Alsace | A1 | 7.544702 | 48.540918 | 18 | 38 | 3.235 | 0.544 | 0.543 | -0.002 | 0.004 |
| A5 | 2021 | Alsace | A1 | 7.564511 | 48.543905 | 26 | 41 | 3.203 | 0.562 | 0.551 | -0.020 | 0.014 |
| A6 | 2021 | Alsace | A1 | 7.571759 | 48.547882 | 30 | 43 | 3.251 | 0.503 | 0.520 | 0.033 | 0.006 |
| A7 | 2021 | Alsace | A1 | 7.575255 | 48.547574 | 33 | 44 | 3.106 | 0.506 | 0.518 | 0.023 | 0.014 |
| A8 | 2022 | Alsace | A1 | 7.579292 | 48.548017 | 36 | 46 | 3.448 | 0.539 | 0.550 | 0.020 | 0.030 |
| A9 | 2021 | Alsace | A1 | 7.586779 | 48.550395 | 31 | 44 | 3.122 | 0.461 | 0.484 | 0.048 | 0.043 |
| BA3 | 2022 | Alsace | A1 | 7.579429 | 48.528832 | 4 | 26 | - | - | - | - | - |
| BA4 | 2022 | Alsace | A1 | 7.581946 | 48.528855 | 3 | 25 | - | - | - | - | - |
| Bli1 | 2022 | Alsace | A4 | 7.490202 | 48.189452 | 30 | 41 | 3.035 | 0.497 | 0.499 | 0.004 | 0.032 |
| Bli2 | 2022 | Alsace | A4 | 7.489811 | 48.218759 | 36 | 44 | 3.102 | 0.519 | 0.496 | -0.046 | 0.020 |
| Bli3 | 2022 | Alsace | A4 | 7.498421 | 48.236530 | 7 | 29 | - | - | - | - | - |
| Bru1 | 2022 | Alsace | A4 | 7.709551 | 48.351107 | 19 | 37 | 3.110 | 0.495 | 0.517 | 0.043 | 0.011 |
| CB2 | 2021 | Alsace | A1 | 7.591170 | 48.554269 | 2 | 19 | - | - | - | - | - |
| CB4 | 2022 | Alsace | A1 | 7.627223 | 48.575197 | 21 | 36 | 3.047 | 0.514 | 0.506 | -0.016 | 0.033 |
| CCR11 | 2022 | Alsace | A4 | 7.721753 | 48.372592 | 27 | 40 | 3.104 | 0.533 | 0.552 | 0.034 | -0.002 |
| CCR12 | 2022 | Alsace | A4 | 7.739052 | 48.346225 | 32 | 40 | 3.207 | 0.547 | 0.549 | 0.004 | 0.015 |
| CCR13 | 2022 | Alsace | A4 | 7.579254 | 48.138813 | 33 | 45 | 3.267 | 0.482 | 0.508 | 0.051 | 0.025 |
| CCR14 | 2022 | Alsace | A4 | 7.726147 | 48.365158 | 21 | 37 | 3.096 | 0.505 | 0.517 | 0.023 | 0.013 |
| CCR2 | 2022 | Alsace | A4 | 7.880296 | 48.660017 | 4 | 28 | - | - | - | - | - |
| CCR4 | 2022 | Alsace | A4 | 7.843604 | 48.646993 | 6 | 29 | - | - | - | - | - |
| CCR7 | 2022 | Alsace | A4 | 7.776558 | 48.500519 | 24 | 37 | 3.024 | 0.508 | 0.522 | 0.027 | 0.034 |
| CCR8 | 2022 | Alsace | A4 | 7.762018 | 48.487490 | 3 | 24 | - | - | - | - | - |
| Ceh0 | 2022 | Alsace | A2 | 7.576856 | 48.475591 | 4 | 28 | - | - | - | - | - |
| Ceh1 | 2021 | Alsace | A2 | 7.581641 | 48.478382 | 31 | 43 | 3.370 | 0.542 | 0.568 | 0.046 | 0.016 |
| Ceh2 | 2022 | Alsace | A2 | 7.588845 | 48.482536 | 27 | 37 | 3.077 | 0.515 | 0.547 | 0.059 | 0.021 |
| CHA1 | 2022 | Alsace | A1 | 7.454961 | 48.573777 | 29 | 44 | 3.351 | 0.559 | 0.544 | -0.028 | 0.014 |
| CHA12 | 2022 | Alsace | A1 | 7.470093 | 48.574551 | 30 | 45 | 3.429 | 0.540 | 0.556 | 0.029 | 0.002 |
| CHA2 | 2022 | Alsace | A1 | 7.479667 | 48.577296 | 30 | 41 | 3.170 | 0.510 | 0.532 | 0.041 | 0.013 |
| Dau1 | 2022 | Alsace | A4 | 7.712416 | 48.374920 | 31 | 43 | 3.305 | 0.561 | 0.550 | -0.020 | 0.008 |
| Deu1 | 2022 | Alsace | A3 | 7.767138 | 48.098375 | 28 | 43 | 3.337 | 0.546 | 0.536 | -0.019 | 0.029 |
| Deu2 | 2022 | Alsace | A3 | 7.760566 | 48.091790 | 2 | 20 | - | - | - | - | - |
| Deu5 | 2022 | Alsace | A3 | 7.868078 | 48.096462 | 23 | 40 | 3.198 | 0.509 | 0.521 | 0.023 | 0.036 |
| Deu7 | 2022 | Alsace | A3 | 7.815028 | 48.045885 | 38 | 50 | 3.347 | 0.511 | 0.531 | 0.038 | 0.032 |
| Deu8 | 2022 | Alsace | A3 | 7.763198 | 48.099368 | 22 | 37 | 2.971 | 0.536 | 0.516 | -0.039 | 0.031 |
| Ehn1 | 2021 | Alsace | A2 | 7.563860 | 48.470101 | 20 | 38 | 3.246 | 0.520 | 0.567 | 0.083 | 0.008 |
| Ehn9 | 2022 | Alsace | A2 | 7.553003 | 48.457075 | 3 | 31 | - | - | - | - | - |
| Eng1 | 2022 | Alsace | A1 | 7.401690 | 48.565821 | 25 | 44 | 3.312 | 0.560 | 0.555 | -0.009 | 0.014 |
| Far1 | 2022 | Alsace | A4 | 7.405380 | 48.278538 | 2 | 20 | - | - | - | - | - |
| Fri1 | 2022 | Alsace | A4 | 7.543433 | 48.291327 | 29 | 42 | 3.231 | 0.521 | 0.523 | 0.004 | 0.015 |
| Fri2 | 2022 | Alsace | A4 | 7.547780 | 48.302849 | 26 | 40 | 3.158 | 0.523 | 0.515 | -0.016 | 0.020 |
| Ger1 | 2022 | Alsace | A4 | 7.710864 | 48.397795 | 33 | 40 | 3.218 | 0.576 | 0.554 | -0.040 | 0.018 |
| Ger3 | 2021 | Alsace | A4 | 7.712247 | 48.388783 | 21 | 41 | 3.332 | 0.500 | 0.533 | 0.062 | 0.007 |
| Ger4 | 2022 | Alsace | A4 | 7.680718 | 48.351555 | 30 | 39 | 3.103 | 0.540 | 0.520 | -0.038 | 0.014 |
| Ger6 | 2021 | Alsace | A4 | 7.674729 | 48.328929 | 20 | 39 | 3.174 | 0.560 | 0.526 | -0.065 | 0.009 |
| Gox1 | 2022 | Alsace | A2 | 7.487110 | 48.434042 | 27 | 34 | 2.919 | 0.533 | 0.499 | -0.068 | 0.124 |
| Han1 | 2021 | Alsace | A4 | 7.699740 | 48.359411 | 25 | 40 | 3.238 | 0.492 | 0.524 | 0.061 | 0.020 |
| Ich0 | 2022 | Alsace | A4 | 7.576146 | 48.188722 | 10 | 35 | 3.309 | 0.590 | 0.544 | -0.085 | 0.021 |
| Jeff1 | 2022 | Alsace | A1 | 7.457562 | 48.577673 | 21 | 42 | 3.356 | 0.543 | 0.513 | -0.058 | 0.041 |
| Kes1 | 2022 | Alsace | A4 | 7.545415 | 48.277385 | 29 | 45 | 3.443 | 0.545 | 0.530 | -0.028 | 0.018 |
| Koh1 | 2022 | Alsace | A1 | 7.421644 | 48.593061 | 34 | 46 | 3.288 | 0.500 | 0.520 | 0.038 | 0.009 |
| Lac1 | 2022 | Alsace | A4 | 7.703560 | 48.376597 | 19 | 41 | 3.174 | 0.537 | 0.533 | -0.008 | 0.023 |
| Lan1 | 2022 | Alsace | A4 | 7.563761 | 48.312189 | 34 | 39 | 3.034 | 0.562 | 0.518 | -0.085 | 0.010 |
| Mul5 | 2022 | Alsace | A1 | 7.608110 | 48.577487 | 6 | 27 | - | - | - | - | - |
| Ost1 | 2022 | Alsace | A4 | 7.706748 | 48.558034 | 4 | 23 | - | - | - | - | - |
| Rla23 | 2021 | Alsace | A6 | 7.668104 | 48.673981 | 3 | 26 | - | - | - | - | - |
| Rla3 | 2021 | Alsace | A6 | 7.680047 | 48.671614 | 20 | 35 | 3.006 | 0.595 | 0.537 | -0.108 | 0.042 |
| Rla4 | 2022 | Alsace | A6 | 7.687878 | 48.671665 | 3 | 23 | - | - | - | - | - |
| Rla5 | 2022 | Alsace | A6 | 7.692171 | 48.671474 | 27 | 45 | 3.320 | 0.552 | 0.550 | -0.004 | 0.010 |
| Rla6 | 2021 | Alsace | A6 | 7.700949 | 48.670594 | 29 | 47 | 3.244 | 0.521 | 0.519 | -0.004 | 0.026 |
| Rlei23 | 2021 | Alsace | A5 | 7.602713 | 48.655405 | 24 | 47 | 3.374 | 0.600 | 0.558 | -0.075 | -0.003 |
| Rlei45 | 2022 | Alsace | A5 | 7.633774 | 48.649009 | 30 | 45 | 3.417 | 0.513 | 0.551 | 0.069 | 0.056 |
| Rlei6 | 2022 | Alsace | A5 | 7.652179 | 48.644755 | 29 | 42 | 3.230 | 0.507 | 0.542 | 0.065 | 0.010 |
| Rlei7 | 2022 | Alsace | A5 | 7.654957 | 48.644172 | 34 | 44 | 3.165 | 0.535 | 0.529 | -0.011 | 0.002 |
| Rlei8 | 2022 | Alsace | A5 | 7.664481 | 48.643140 | 26 | 45 | 3.159 | 0.538 | 0.527 | -0.021 | 0.003 |
| RLK1 | 2021 | Alsace | A5 | 7.635142 | 48.658617 | 23 | 41 | 3.156 | 0.565 | 0.522 | -0.082 | 0.002 |
| RLK2 | 2022 | Alsace | A5 | 7.651268 | 48.657640 | 33 | 43 | 3.176 | 0.542 | 0.540 | -0.004 | 0.010 |
| RLK3 | 2022 | Alsace | A5 | 7.660315 | 48.656235 | 15 | 38 | 3.245 | 0.513 | 0.552 | 0.071 | -0.001 |
| RLK4 | 2022 | Alsace | A5 | 7.664291 | 48.654240 | 23 | 40 | 3.191 | 0.587 | 0.536 | -0.095 | 0.027 |
| RLK56 | 2022 | Alsace | A5 | 7.678233 | 48.650551 | 18 | 36 | 3.036 | 0.528 | 0.530 | 0.004 | 0.037 |
| Rlmu1 | 2022 | Alsace | A5 | 7.614947 | 48.608272 | 3 | 28 | - | - | - | - | - |
| Rlmu2 | 2022 | Alsace | A5 | 7.626599 | 48.607025 | 20 | 37 | 2.980 | 0.525 | 0.482 | -0.089 | 0.015 |
| Rlmu3 | 2022 | Alsace | A5 | 7.629157 | 48.607368 | 4 | 26 | - | - | - | - | - |
| Rmh1 | 2021 | Alsace | A6 | 7.688742 | 48.685199 | 7 | 32 | - | - | - | - | - |
| Rsc1 | 2022 | Alsace | A4 | 7.739234 | 48.482301 | 20 | 40 | 3.305 | 0.520 | 0.532 | 0.023 | 0.061 |
| Rsc2 | 2021 | Alsace | A4 | 7.741570 | 48.490704 | 1 | 19 | - | - | - | - | - |
| Rsc3 | 2021 | Alsace | A4 | 7.737795 | 48.493810 | 3 | 28 | - | - | - | - | - |
| Rut2 | 2022 | Alsace | A2 | 7.595870 | 48.499645 | 28 | 38 | 3.144 | 0.525 | 0.523 | -0.004 | 0.040 |
| Rut3 | 2022 | Alsace | A2 | 7.605300 | 48.500283 | 23 | 39 | 3.314 | 0.561 | 0.586 | 0.043 | 0.010 |
| Rut4 | 2022 | Alsace | A2 | 7.612676 | 48.501593 | 9 | 34 | 3.320 | 0.589 | 0.544 | -0.083 | 0.036 |
| S12 | 2022 | Alsace | A5 | 7.688485 | 48.632729 | 5 | 26 | - | - | - | - | - |
| S17 | 2021 | Alsace | A5 | 7.741153 | 48.641593 | 21 | 38 | 3.043 | 0.476 | 0.508 | 0.063 | 0.110 |
| S21 | 2021 | Alsace | A4 | 7.833577 | 48.633176 | 25 | 39 | 3.339 | 0.556 | 0.567 | 0.019 | 0.077 |
| S2 | 2022 | Alsace | A5 | 7.562404 | 48.638451 | 26 | 43 | 3.308 | 0.500 | 0.539 | 0.072 | 0.002 |
| S3 | 2021 | Alsace | A5 | 7.570723 | 48.634705 | 34 | 43 | 3.201 | 0.526 | 0.534 | 0.015 | 0.009 |
| S6 | 2021 | Alsace | A5 | 7.615283 | 48.629329 | 31 | 42 | 3.176 | 0.516 | 0.519 | 0.006 | 0.005 |
| S67 | 2022 | Alsace | A5 | 7.622262 | 48.630331 | 20 | 38 | 3.089 | 0.475 | 0.517 | 0.081 | 0.000 |
| S677 | 2022 | Alsace | A5 | 7.629693 | 48.630449 | 4 | 27 | - | - | - | - | - |
| Sci1 | 2022 | Alsace | A4 | 7.541203 | 48.257234 | 2 | 21 | - | - | - | - | - |
| Ser1 | 2022 | Alsace | A2 | 7.436113 | 48.344174 | 4 | 25 | - | - | - | - | - |
| V1 | 2021 | Alsace | A4 | 7.607497 | 48.192280 | 35 | 45 | 3.263 | 0.554 | 0.537 | -0.032 | 0.004 |
| V2 | 2022 | Alsace | A4 | 7.674399 | 48.278712 | 30 | 36 | 2.958 | 0.500 | 0.504 | 0.008 | 0.012 |
| V3 | 2022 | Alsace | A4 | 7.732873 | 48.358055 | 33 | 40 | 3.158 | 0.536 | 0.548 | 0.022 | 0.002 |
| V4 | 2022 | Alsace | A4 | 7.719338 | 48.325092 | 34 | 45 | 3.161 | 0.518 | 0.527 | 0.017 | 0.010 |
| Wei1 | 2022 | Alsace | A4 | 7.763958 | 48.494377 | 27 | 37 | 3.040 | 0.474 | 0.508 | 0.067 | 0.029 |
| Wur1 | 2022 | Alsace | A4 | 7.705175 | 48.351641 | 32 | 44 | 3.245 | 0.541 | 0.530 | -0.021 | 0.029 |
| Zp0 | 2022 | Alsace | A6 | 7.491934 | 48.742628 | 35 | 40 | 2.982 | 0.514 | 0.494 | -0.040 | 0.033 |
| Zp1 | 2022 | Alsace | A6 | 7.499699 | 48.742984 | 30 | 40 | 3.075 | 0.500 | 0.500 | 0.000 | 0.047 |
| Zp2 | 2022 | Alsace | A6 | 7.553629 | 48.751473 | 33 | 41 | 3.042 | 0.461 | 0.500 | 0.078 | 0.029 |
| Bai1 | 2017 | Nord | N3 | 1.814731 | 50.556417 | 34 | 38 | 3.102 | 0.502 | 0.484 | -0.037 | 0.082 |
| Bai2 | 2017 | Nord | N3 | 1.795901 | 50.547208 | 28 | 40 | 3.223 | 0.504 | 0.501 | -0.006 | 0.067 |
| Bai3 | 2017 | Nord | N3 | 1.784821 | 50.531367 | 32 | 32 | 2.538 | 0.373 | 0.389 | 0.041 | 0.148 |
| Bai4 | 2017 | Nord | N3 | 1.786619 | 50.521859 | 4 | 23 | - | - | - | - | - |
| Bam1 | 2017 | Nord | N6 | 1.743292 | 49.241668 | 31 | 34 | 2.868 | 0.506 | 0.493 | -0.026 | 0.059 |
| Bav1 | 2017 | Nord | N6 | 1.729459 | 49.242210 | 8 | 32 | 3.200 | 0.575 | 0.563 | -0.021 | 0.129 |
| Cam1 | 2017 | Nord | N3 | 1.606164 | 50.573788 | 36 | 41 | 3.321 | 0.503 | 0.511 | 0.016 | 0.028 |
| Char1 | 2017 | Nord | N2 | 2.605009 | 49.128569 | 33 | 43 | 3.403 | 0.552 | 0.528 | -0.045 | 0.060 |
| Cuy | 2022 | Nord | N4 | 3.458695 | 50.442091 | 20 | 31 | 2.623 | 0.435 | 0.418 | -0.041 | 0.226 |
| Foc1 | 2018 | Nord | N1 | 5.072654 | 50.129394 | 19 | 39 | 3.311 | 0.500 | 0.537 | 0.069 | 0.047 |
| Fre1 | 2019 | Nord | N5 | 2.067305 | 49.755982 | 26 | 26 | 2.377 | 0.438 | 0.428 | -0.023 | 0.306 |
| Gam1 | 2017 | Nord | N5 | 1.561604 | 49.981019 | 6 | 28 | - | - | - | - | - |
| Gen1 | 2017 | Nord | N1 | 5.115321 | 50.124713 | 17 | 39 | 3.299 | 0.547 | 0.515 | -0.062 | 0.079 |
| Gmar1 | 2017 | Nord | N2 | 2.562805 | 49.137762 | 30 | 40 | 3.326 | 0.520 | 0.533 | 0.024 | 0.034 |
| Hon1 | 2017 | Nord | N3 | 1.611199 | 50.758635 | 33 | 21 | 1.865 | 0.288 | 0.284 | -0.014 | 0.238 |
| Isq | 2017 | Nord | N3 | 1.660956 | 50.668245 | 30 | 35 | 2.913 | 0.434 | 0.468 | 0.073 | 0.128 |
| LGM1 | 2017 | Nord | N5 | 1.870529 | 50.059129 | 29 | 26 | 2.493 | 0.424 | 0.447 | 0.051 | 0.224 |
| MD1 | 2017 | Nord | N5 | 1.862351 | 50.061888 | 15 | 27 | 2.558 | 0.401 | 0.448 | 0.105 | 0.209 |
| Mou1 | 2017 | Nord | N6 | 1.893308 | 49.265562 | 28 | 40 | 3.204 | 0.550 | 0.535 | -0.028 | 0.115 |
| OM1 | 2017 | Nord | N5 | 1.464563 | 50.036032 | 6 | 30 | - | - | - | - | - |
| Rev1 | 2017 | Nord | N6 | 1.871270 | 49.226970 | 9 | 29 | 2.853 | 0.467 | 0.500 | 0.066 | 0.193 |
| SMar1 | 2017 | Nord | N2 | 2.461970 | 49.150454 | 8 | 32 | 3.200 | 0.550 | 0.587 | 0.063 | 0.142 |
| Tou1 | 2017 | Nord | N2 | 2.590224 | 49.127125 | 31 | 40 | 3.157 | 0.520 | 0.533 | 0.024 | 0.074 |
| Wal1 | 2017 | Nord | N4 | 3.373399 | 50.389700 | 32 | 24 | 2.271 | 0.462 | 0.429 | -0.077 | 0.264 |

**Table S2:** Estimates of genetic diversity for each microsatellite locus. Multiplex: multiplex in which the microsatellite locus was amplified, Fluo: fluorophore used for genotyping (FAM: blue, NED: yellow, PET: red, VIC: green), *A*N: number of alleles, *H*o: observed heterozygosity, *H*e: expected heterozygosity, *F*IS: intrapopulation fixation index, *F*ST: mean genetic differentiation across sampled populations. *: *P <* 0.05, ***: *P* < 0.001.

| Locus | Multiplex | Fluo | $A_N$ | $H_o$ | $H_e$ | $F_{ST}$ | $F_{IS}$ |
| --- | --- | --- | --- | --- | --- | --- | --- |
| LIST024 | PLEX1 | VIC | 5 | 0.643 | 0.666 | 0.035 *** | -0.001 |
| LIST034 | PLEX1 | PET | 2 | 0.471 | 0.500 | 0.073 *** | -0.008 |
| LIST035 | PLEX1 | VIC | 24 | 0.652 | 0.706 | 0.070 *** | 0.006 |
| LIST063 | PLEX1 | FAM | 12 | 0.629 | 0.673 | 0.061 *** | -0.001 |
| LIST066 | PLEX1 | NED | 14 | 0.544 | 0.583 | 0.069 *** | -0.004 |
| LIST002 | PLEX2 | VIC | 2 | 0.367 | 0.401 | 0.074 *** | -0.012 |
| LIST023 | PLEX2 | VIC | 11 | 0.639 | 0.687 | 0.054 *** | 0.004 |
| LIST037 | PLEX2 | FAM | 11 | 0.320 | 0.347 | 0.062 *** | 0.035 * |
| LIST042 | PLEX2 | VIC | 5 | 0.591 | 0.655 | 0.081 *** | 0.010 |
| LIST060 | PLEX2 | NED | 6 | 0.411 | 0.689 | 0.053 *** | 0.377 *** |
| LIST062 | PLEX2 | PET | 4 | 0.311 | 0.335 | 0.088 *** | -0.005 |
| Mean over all loci |  |  |  |  |  | 0.063 * | 0.045 |
| Mean over all loci without locus LIST060 |  |  |  |  |  | 0.065 * | 0.002 |

**Table S3:**
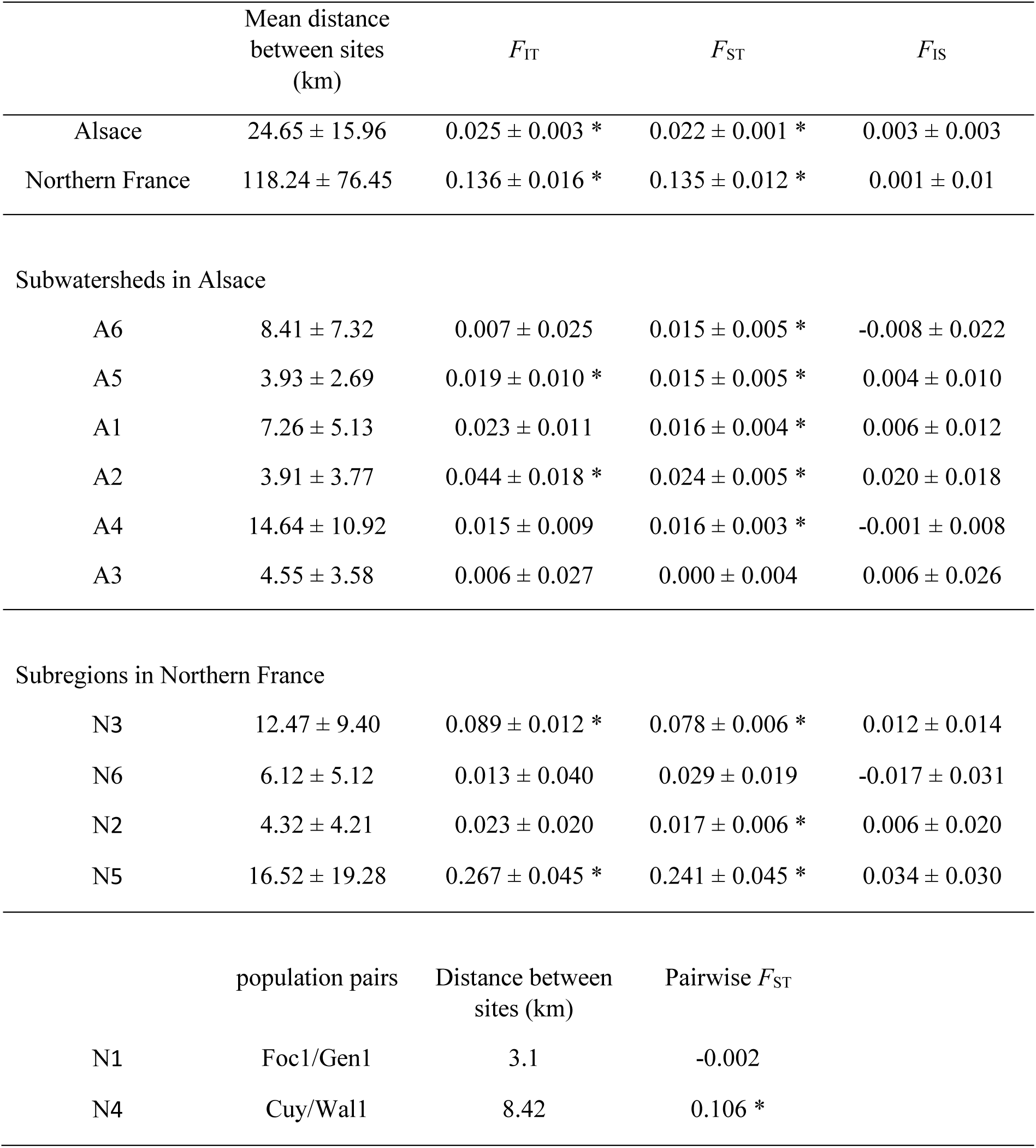
Weir and Cockerham’s (1984) estimates of *F*-statistics for Alsace and northern France region and subregions or subwatersheds. In the two subregions where only two populations were sampled, only the pairwise *F*ST is reported. *: *P <* 0.05.

